# Investigating β-lactam drug targets in *Mycobacterium tuberculosis* using chemical probes

**DOI:** 10.1101/2019.12.19.881631

**Authors:** Samantha R. Levine, Kimberly E. Beatty

## Abstract

Tuberculosis is a deadly disease that requires a flexible arsenal of drugs to treat it. Although β-lactam antibiotics are rarely used to treat *Mycobacterium tuberculosis* (*Mtb*) infections today, the targets of these drugs are present in the bacterium. Moreover, the cell wall peptidoglycan of *Mtb* contains an abundance of unusual (3→3) cross-links. These cross-links are formed by enzymes called L,D-transpeptidases, which are susceptible to inhibition by the carbapenem class of antibiotics. We developed new small molecule probes to investigate the L,D-transpeptidases and other β-lactam drug targets in *Mtb*. We synthesized probes based on three classes of antibiotics, a monobactam, cephalosporin, and carbapenem. For the carbapenem, we synthesized a meropenem analogue conjugated to a far-red fluorophore. This probe was particularly useful in identifying active L,D-transpeptidases in protein gel-resolved lysates. Next we analyzed β-lactam targets in lysates from both hypoxic and actively-replicating cultures of *Mtb*. We identified numerous targets, including transpeptidases, carboxypeptidases, and the β-lactamase BlaC. Overall, we provide evidence that *Mtb* dynamically regulates the enzymes responsible for maintaining cell wall peptidoglycan and that meropenem is a good inhibitor of those enzymes.

## Introduction

Tuberculosis (TB) is the most deadly infectious disease in human history. Over time, *Mycobacterium tuberculosis* (*Mtb*) strains have developed resistances to all anti-mycobacterial drugs. Drug-resistant strains occur worldwide and cause ~6% of TB infections^*(1)*^. The global rise of drug resistance necessitates finding new drugs, a slow and expensive process, or repurposing existing drugs^*(2–4)*^. New TB treatments would ideally target both active TB and latent TB infections (LTBI). In the human host, *Mtb* encounters environmental stresses that can trigger a dormant, non-replicating state. These dormant bacteria exhibit phenotypic drug resistance to most front-line drugs^*(5–7)*^. While dormant *Mtb* is associated with LTBIs, active TB also presents with metabolically diverse populations^*(5–7)*^. There is an urgent need to identify and validate therapeutics that will be broadly useful for the majority of TB patients.

β-lactam antibiotics are widely used drugs that target cell wall biosynthesis. Two of the earliest classes of β-lactams, the penicillins and cephalosporins, inhibit D,D-transpeptidases. These enzymes are penicillin-binding proteins (**PBPs**) that modify cell wall peptidoglycan. They form 4→3 cross-links between a stem peptide’s fourth amino acid, D-Alanine, and the third amino acid on an adjacent stem: meso-diaminopimelic acid (**mDAP**)^35,36^. In most bacteria, the peptidoglycan is comprised of 4→3 linkages. However, in *Mtb* 3→3 cross-links predominate and comprise up to 80% of linkages*^(8, 9)^*. This covalent modification, from mDAP^3^ to mDAP^3^, is generated by L,D-transpeptidases (**LDTs**). A recent review describes the structure and function of *Mtb*’s PBPs and LDTs^*(10)*^. Notably, Ldt_Mt1_ and Ldt_Mt2_ are essential for virulence*^(11, 12)^*, and Ldt_Mt1_ is believed to remodel the cell wall for survival in dormancy*^(8, 13)^*.

LDTs and PBPs have different β-lactam susceptibilities, as evidenced by enzymatic, biochemical, and structural studies^*(14–20)*^. For example, PBPs are inactivated by covalent modification at a catalytic serine by β-lactam antibiotics (e.g., penams, monobactams, cephems, and carbapenems). In contrast, LDTs are inhibited by compounds in the carbapenem class, which bind to an active-site cysteine by thioester bond formation (Figure 1). Since the majority of cross-links in *Mtb* are formed by LDTs, this suggests that carbapenems could be effective drugs for treating TB*^(10, 21, 22)^*. The *Mtb* LDTs and PBPs are summarized in Table 1.

**Table 1.**
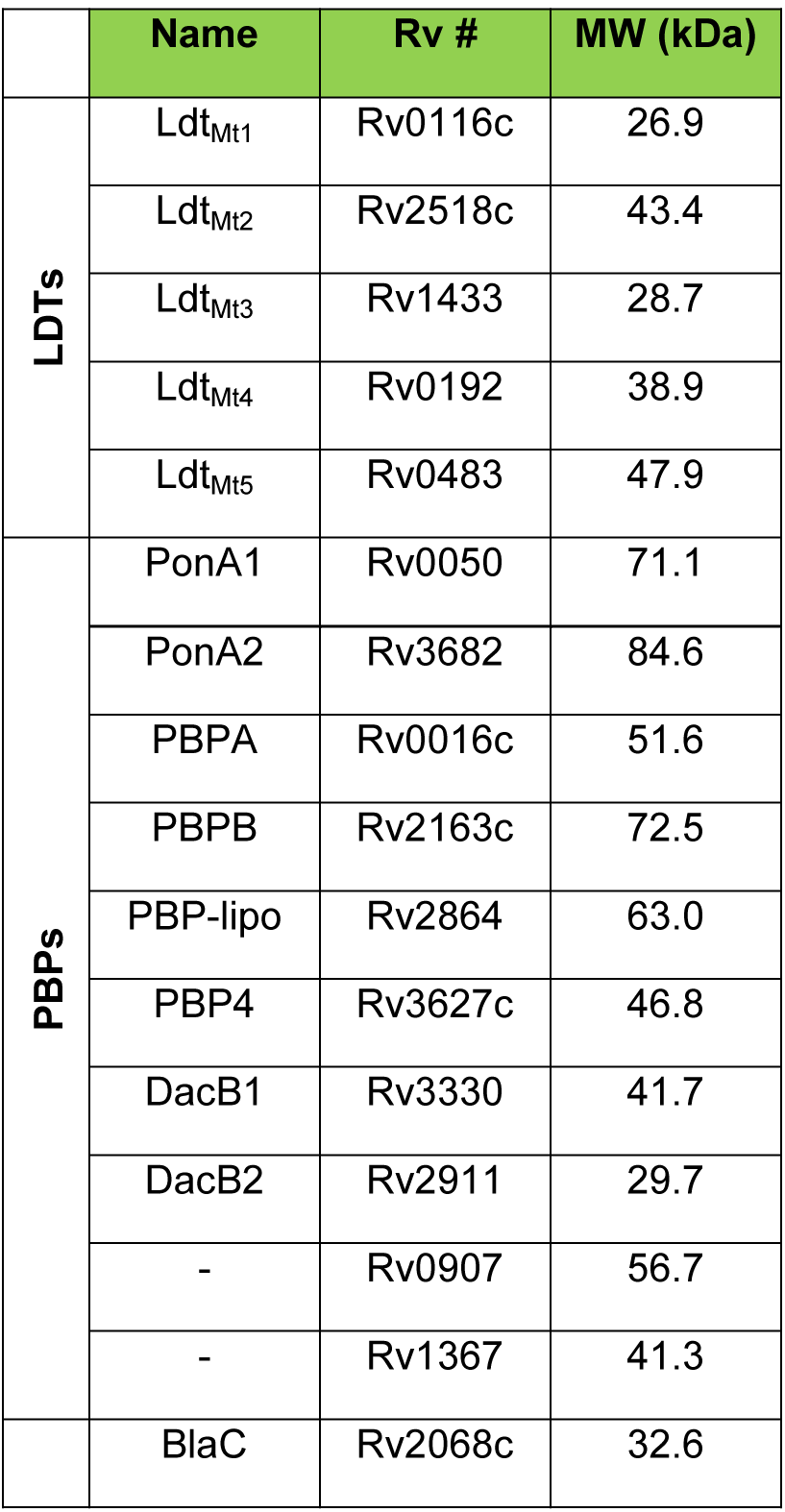
β-lactam targets in *Mtb*.

**Figure 1.**
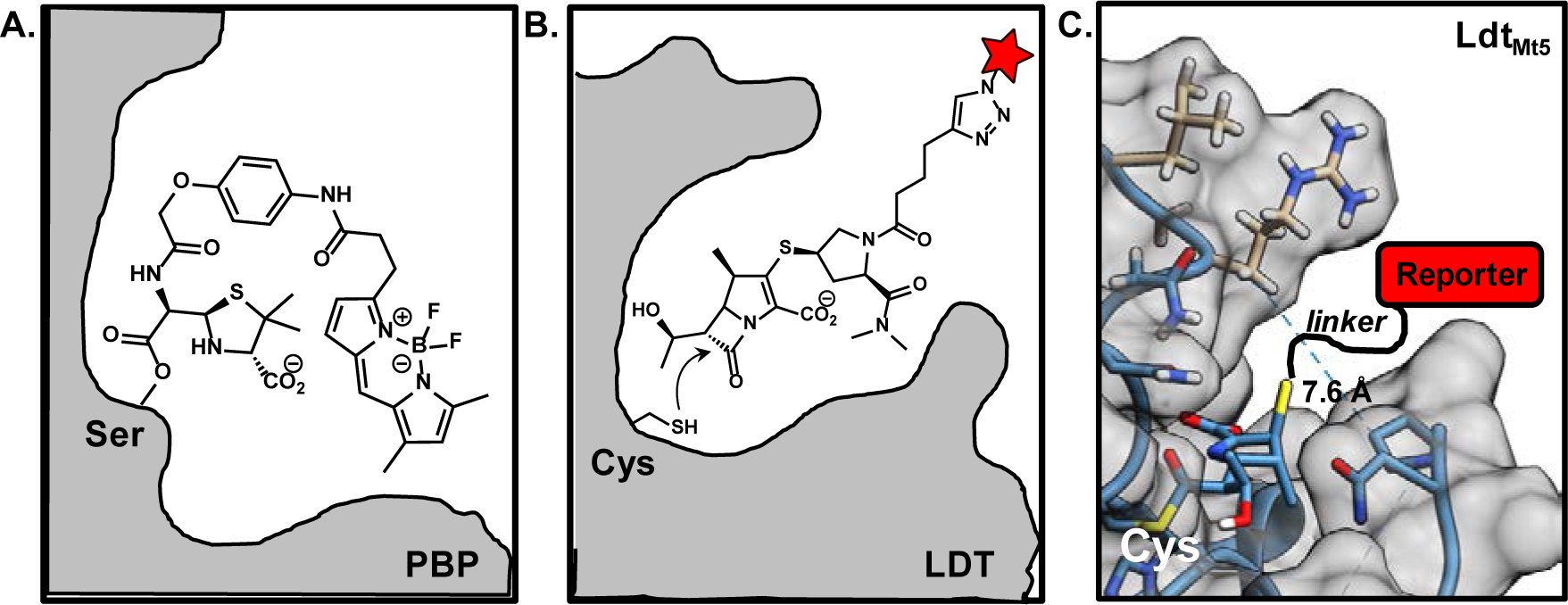
Active site labeling of PBPs and LDTs with β-lactam probes. **A.** In PBPs, the active-site serine binds to a reactive β-lactam probe, as shown for Bocillin FL. **B.** In contrast, the LDT’s active-site cysteine binds to carbapenems (e.g., Mero-Cy5). **C.** Structure of the Ldt_Mt5_ active-site bound to a fragment of meropenem (PDB ID: 4ZFQ)^*(14)*^. The Mero-Cy5 probe, described herein, was designed to bind to LDTs with a reporter extending out of the active site, as illustrated.

β-lactams are not part of the standard treatment regimen for TB. However, the reason for this might lie in outdated observations made 70 years ago, shortly after the introduction of penicillin. A study from 1949 described a “penicillinase” that made *Mtb* resistant to β-lactam antibiotics^*(23)*^. This resistance was conferred by BlaC, a genomically-encoded β-lactamase*^(24, 25)^*. The discovery of BlaC suggested that β-lactam antibiotics would be ineffective in treating TB. BlaC also inactivated the cephalosporins introduced in the 1960s^*(26)*^. Yet, as reviewed by Story-Roller and Lamichhane^*(22)*^, the carbapenem class targets LDTs, which may provide distinct druggable targets in *Mtb*. BlaC has limited activity against carbapenems^*(27–29)*^.

There is some evidence that β-lactams are therapeutically-relevant drugs for TB^*(22)*^. There have been occasional reports of successfully treating TB patients with co-administration of β-lactams with clavulanate, a BlaC inhibitor*^(21, 30–36)^*. A 1998 report described using amoxicillin/clavulanate to cure multi-drug resistant TB^*(35)*^. Drug-resistant TB has also been treated with imipenem*^(37)^* and meropenem/clavulanate*^(30–32, 34, 36)^*. The latter treatment cured extensively-drug resistant TB patients (83% cure rate)^*(34)*^. This combination was also effective against dormant *Mtb^(38, 39)^*.

One outcome of the prior studies is a renewed interest in identifying β-lactams that can cure drug-resistant TB. The scarcity of information on their protein targets in *Mtb* makes the development of broadly successful treatment regimens problematic. Furthermore, there is almost no information on which PBPs and LDTs are functional in dormant and actively replicating *Mtb*. Prior transcriptional analyses suggest that these enzymes are dynamically regulated by the pathogen based on its metabolic state*^(13, 40)^*. However, transcript levels do not always correlate with enzyme activity.

Evaluating multiple enzymes at once is challenging using conventional biochemical or genetic approaches. Activity-based probes (ABPs) enable such studies. ABPs consist of a reporter linked to an irreversible inhibitor, which provides specificity for the enzyme target^*(41–43)*^. There is established precedence for using ABPs to assign peptidase functions in other bacteria^*(44–49)*^.

In this work, we used ABPs to profile the regulation of *Mtb* PBPs and LDTs. We produced a small set of structurally-diverse β-lactam probes, including a new meropenem derivative, to illuminate drug-bound targets in protein gel-resolved mycobacterial lysates. We identified enzymes associated with both dormant and actively-replicating *Mtb*. Our results suggest that carbapenems, including meropenem, could be useful drugs for treating latent or active TB infections.

## Results and Discussion

### Identification of β-lactam targets encoded in the *Mtb* genome

We initiated this project by generating a comprehensive list of *Mtb* enzymes with active sites likely to interact with a β-lactam. We searched the *Mtb* H37Rv genome for genes with protein family (i.e., Pfam) annotation indicating potential PBP, LDT, or β-lactamase activity.*^(50)^* This gave a total of 43 proteins with either validated or theoretical abilities to interact with β-lactams (see *Supporting Information*, **Table S1**). The abbreviated version of this list (Table 1) contains only the most likely β-lactam targets, and includes ten PBPs, five LDTs, and the β-lactamase BlaC^*(10)*^.

### Synthesis and validation of fluorescent β-lactam probes

Next, we assembled our chemical toolkit for detecting and identifying β-lactam targets in *Mtb* (Figure 2). For our studies, all ABPs contained a β-lactam antibiotic as the inhibitor. For a penam compound we obtained Bocillin-FL, a commercially-available green fluorescent penicillin^*(44)*^. We additionally selected monobactam, cephalosporin, and carbapenem scaffolds for structural diversity. An alkyne handle was included in these ABPs to enable ready access to fluorescent probes via azide-alkyne conjugation.

**Figure 2.**
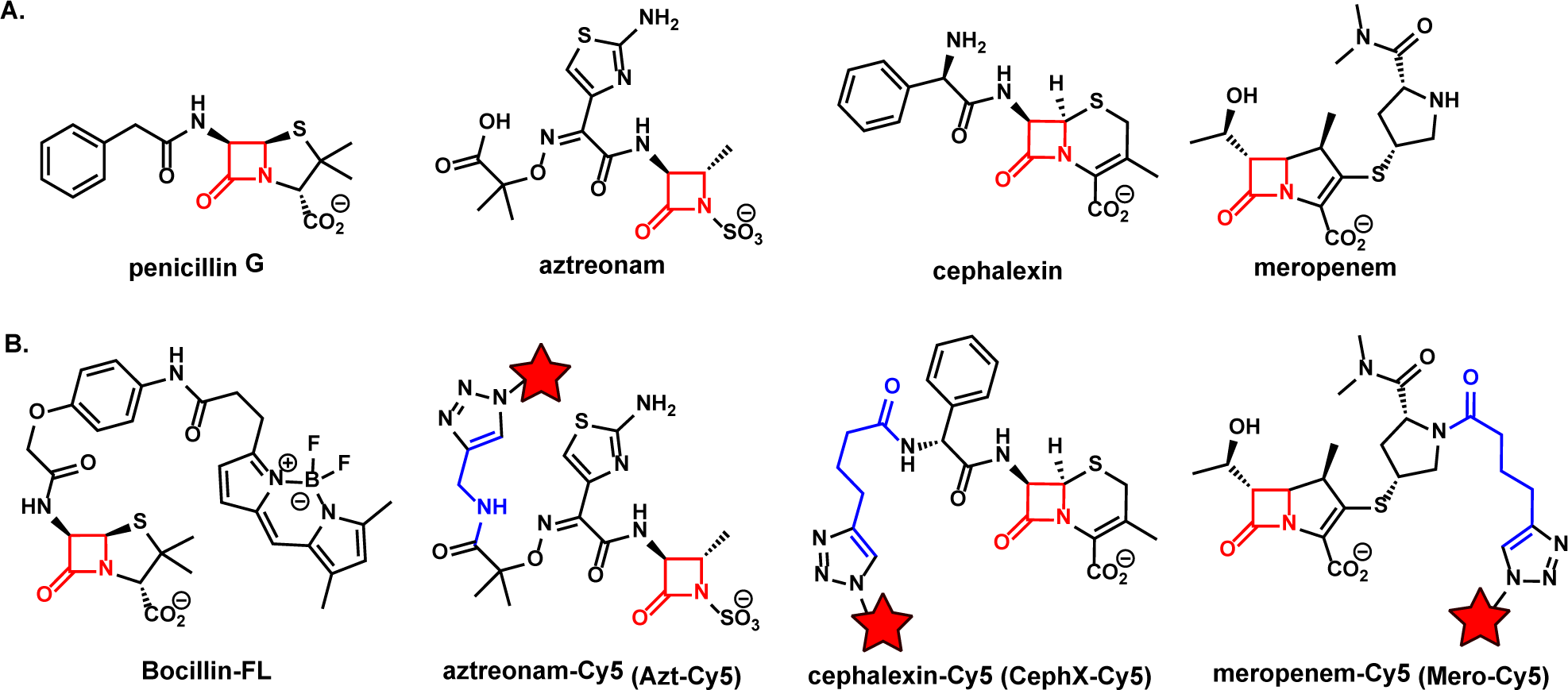
Structures of β-lactam antibiotics (A) and activity-based probes (B) used in the current work. The Cy5-conjugated derivatives were obtained by reacting the alkyne probe with sulfo-Cy5-azide to produce Azt-Cy5, CephX-Cy5, and Mero-Cy5. The red star signifies the sulfo-Cy5 fluorophore.

For the monobactam compound, we synthesized the known aztreonam-N-alkyne^*(48)*^. We found that increasing the reaction time resulted in a substantially higher yield of aztreonam-alkyne than previously reported (>95% yield in 3 d vs. 19% in 30 min).^*(48)*^ For the cephalosporin and carbapenem scaffolds we synthesized two new compounds: cephalexin-alkyne and meropenem-alkyne, respectively. For the cephalosporin probe, we used the readily available cephalexin as the starting material, providing a cost benefit over the more traditional cephalosporin C.*^(46, 48)^* We synthesized cephalexin-alkyne under basic conditions similar to ones used by Carlson and co-workers to synthesize cephalosporin C probes.^*(46)*^ A modification of these conditions to use a heterogeneous base (Cs_2_CO_3_) generated meropenem-alkyne from meropenem. We selected meropenem as our carbapenem-class probe because of its clinical significance for treating TB. Also, it was straightforward to modify, unlike faropenem and imipenem. These alkynes readily underwent copper-catalyzed click reactions with Sulfo-Cyanine5 (Cy5) azide to generate far-red fluorescent probes: aztreonam-Cy5 (Azt-Cy5), cephalexin-Cy5 (CephX-Cy5), and meropenem-Cy5 (Mero-Cy5).

We used Bocillin-FL for our initial evaluation of activity in *Mtb* lysates (Figure 3). Prior work used this probe to detect PBPs—including PonA2—in *Mycobacterium smegmatis* (***Msmeg***), a rapidly growing lab strain of mycobacteria.*^(51, 52)^* We used Bocillin FL to analyze lysates prepared from mid-log phase cultures of *Mtb* (mc^2^6020), a Δ*lysA* Δ*panCD* auxotroph of *Mtb* H37Rv^*(53)*^. Labeled lysates were resolved by SDS-PAGE and imaged on a fluorescence scanner to detect green fluorescence from the target-bound probe (Figure 3). Labeled proteins were nearly all over 50 kDa, despite ten of the targets in Table 1 being of a lower molecular weight. Labeling was reduced by pre-treatment with penicillin G and other β-lactam antibiotics (see *Supplementary Information*; **Figure S1**). This competition experiment demonstrated that labeling was specific for proteins that bind to β-lactams. The fluorescent band intensity corresponded with activity, not relative abundance (see **Figure S1B**). We observed three auto-fluorescent bands in the green channel. Additionally, there were other drawbacks to using Bocillin-FL: (1) Bocillin-FL labeled less than half of the potential targets; (2) it labeled only two proteins under 50 kDa; (3) penicillins are poor inhibitors of LDTs.

**Figure 3.**
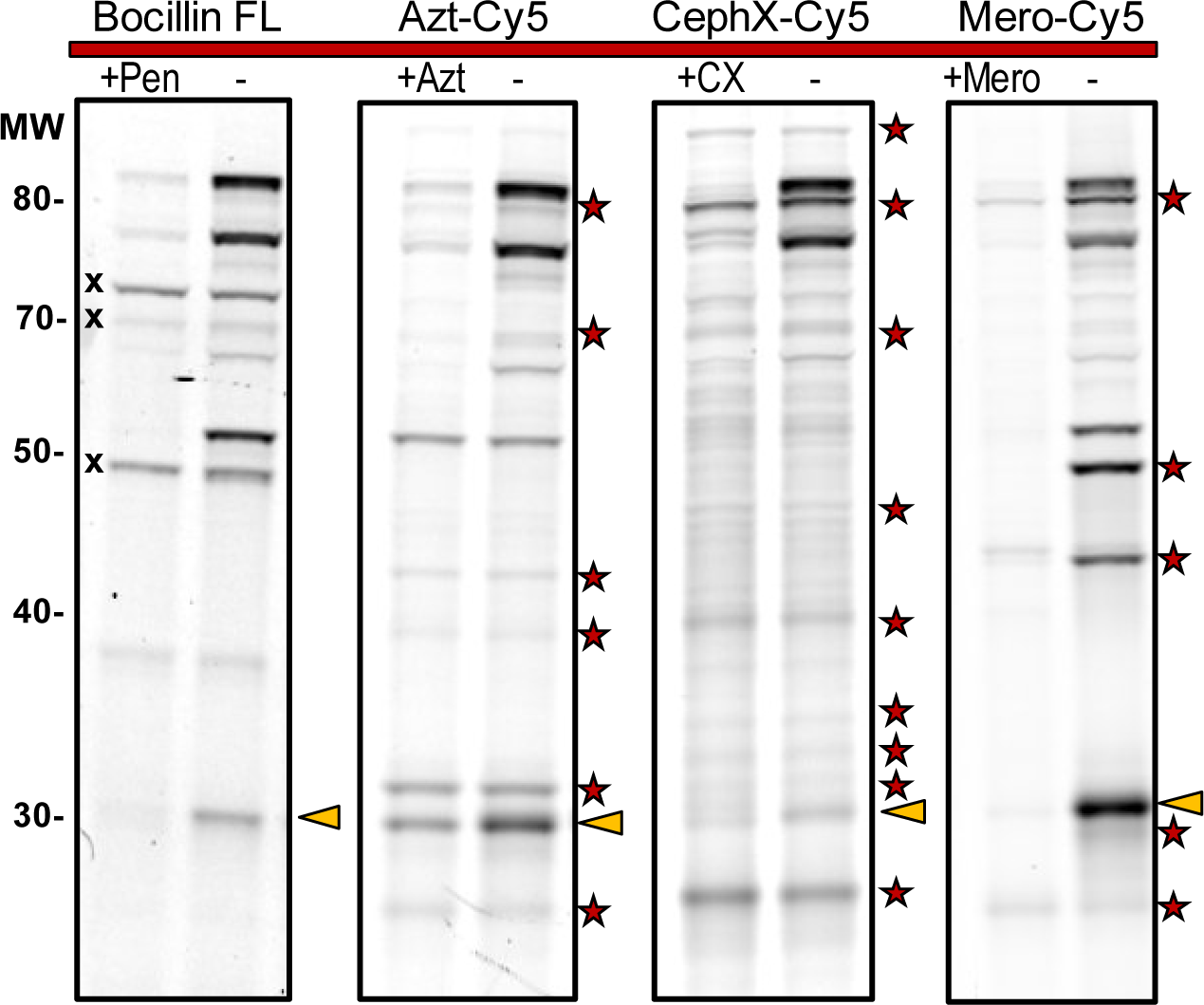
β-lactam probes covalently bind to *Mtb* protein targets, enabling fluorescent detection. Lysates were treated with probe, resolved by SDS-PAGE, and imaged. Three auto-fluorescent bands (x) were observed with Bocillin FL, but not with Cy5 probes. Labeling was reduced by pre-treatment with antibiotic (left lanes). Pen: Penicillin G; Azt: Aztreonam; CX: Cephalexin; Mero: Meropenem. 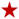: Targets revealed by Azt-Cy5, CephX-Cy5,or Mero-Cy5 (but not Bocillin FL). BlaC activity: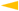.

We overcame those drawbacks by using our far-red fluorescent ABPs: Azt-Cy5, CephX-Cy5, and Mero-Cy5. Again, *Mtb* lysates were probe-treated, resolved by SDS-PAGE, and imaged to detect Cy5-labeled targets (Figure 3). Compared with Bocillin-FL, each of these probes showed more fluorescent bands. They also labeled multiple proteins with an apparent molecular weight of less than 50 kDa. We performed competition experiments with the parent antibiotic. Labeling was reduced under these conditions, indicating that the labeling was specific for proteins that bind β-lactams. Notably, these new probes each revealed distinct target selectivity plus bands that were undetected by Bocillin-FL. All four probes detected BlaC activity (see next section).

### Identification of protein targets of Mero-Cy5 from gel-excised bands

We initiated the process of identifying proteins that were labeled by Mero-Cy5 because this compound provided the clearest in-gel activity patterns. In these studies we compared lysates from actively-replicating cultures with dormant cultures of *Mtb*. We induced dormancy by incubation under hypoxic conditions, as described^*(54–56)*^. *Mtb* lysates were fractionated by centrifugation (into pellet and supernatant^*(57)*^), labeled with Mero-Cy5, and resolved by SDS-PAGE. Fluorescent bands were excised and submitted for identification by liquid chromatography tandem mass spectrometry (LC-MS/MS). Peptides were considered positively identified if their probability of identification was ≥99% with a false-discovery rate (FDR) of 0.67%. Proteins were considered positively identified if two or more exclusive unique peptides were detected (probability of identification ≥95%; FDR 0.0%).

We narrowed our analysis to proteins with putative PBP, LDT, or β-lactamase activity (see **Table S1**). Of the forty-three proteins in **Table S1**, thirty-two of them were identified in our samples (**Table S2**). Specifically, we identified transpeptidases (PonA1, PonA2, PBPA, LDT_Mt2_, LDT_Mt3_, LDT_Mt5_), carboxypeptidases (DacB1, DacB2, PBP4), and the β-lactamase BlaC. The remaining proteins detected were confirmed or putative β-lactamases based on Pfam annotation. We attempted to precisely extract bands based on fluorescent signal, but the presence of a particular protein does not indicate that it was a source of the Cy5 signal in a given region. Experimental details and the full data set are provided in the SI, including annotated gels from which we excised bands (**Figure S2**).

Band identifications are provided in Figures 4–6. For clarity, bands were divided into three regions based on migration and molecular weight: high (~80 kDa), middle (40-50 kDa), and low (15-30 kDa), as indicated in Figure 4A. First we analyzed bands that migrated at or below 30 kDa. We determined that BlaC activity most likely contributes much of the signal at 30 kDa in normoxic samples (Band ID: 10NP8). We identified 18 unique peptides from BlaC with 69.7% sequence coverage. We validated our assignment by comparing lysates from *Mtb* mc^2^6020 with *Mycobacterium marinum*, a close genetic relative, and with *Msmeg* (mc^2^155), a species that lacks a BlaC ortholog (**Figure S3**). The major β-lactamase in *Msmeg*, BlaS (31 kDa), is not a target of this inhibitor*^(24, 58, 59)^*. We observed a clavulanate-sensitive band at 30 kDa for lysates from *Mtb* and *M. marinum*, but not *Msmeg*. This comparative analysis provides compelling evidence in support of this assignment.

**Figure 4.**
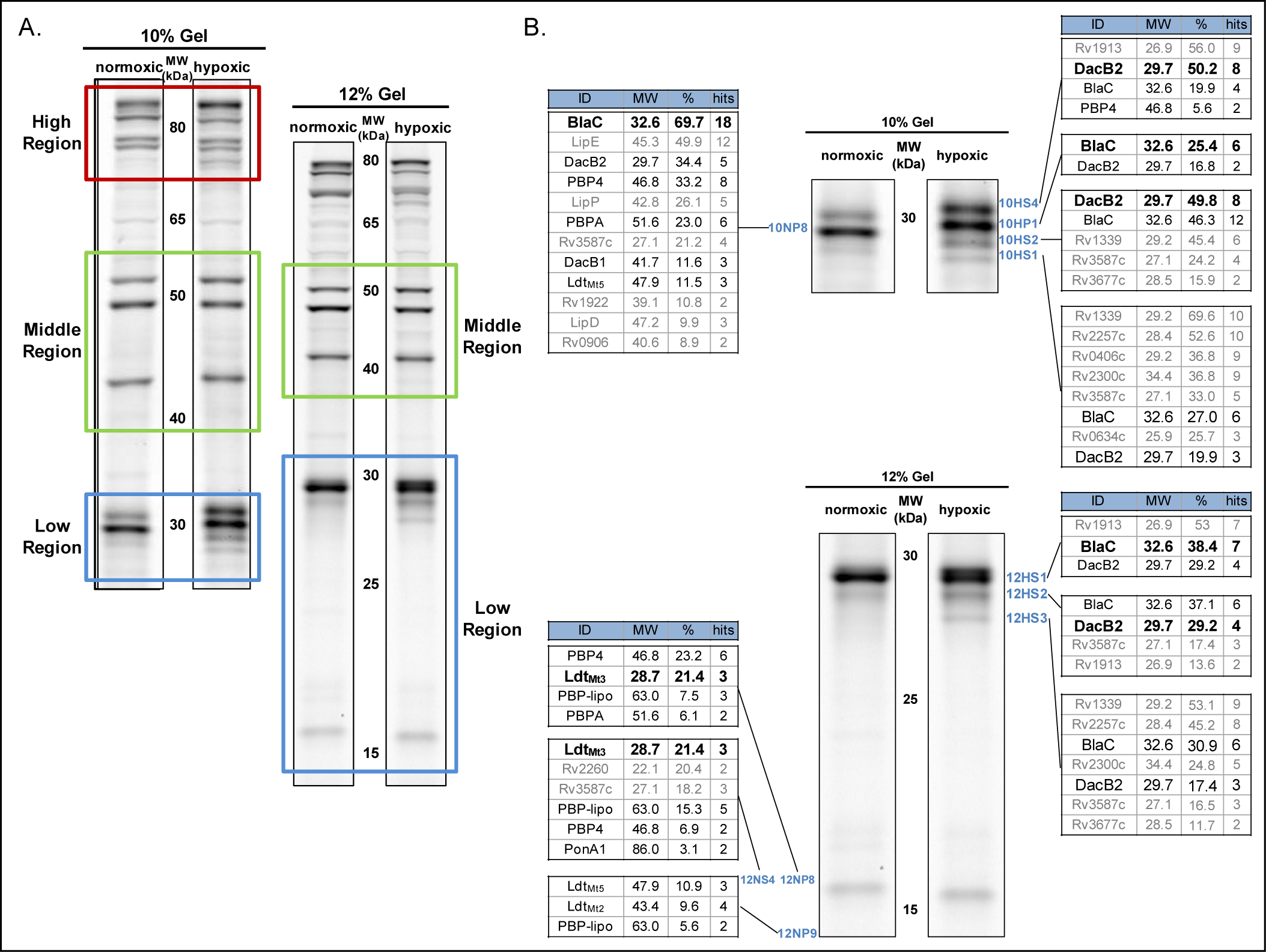
Analysis of lysates for MS-based identification of proteins labeled with Mero-Cy5. **A.** Visualization of the division of protein gels into High, Middle, and Low regions. **B.** Protein identifications for bands excised from the “low region” of the 10% (top) and 12% (bottom) SDS-PAGE gels. The most likely source of the fluorescent signal observed is emphasized in **Bold**. Known targets of β-lactams are indicated in black, while putative targets are in gray.

**Figure 5.**
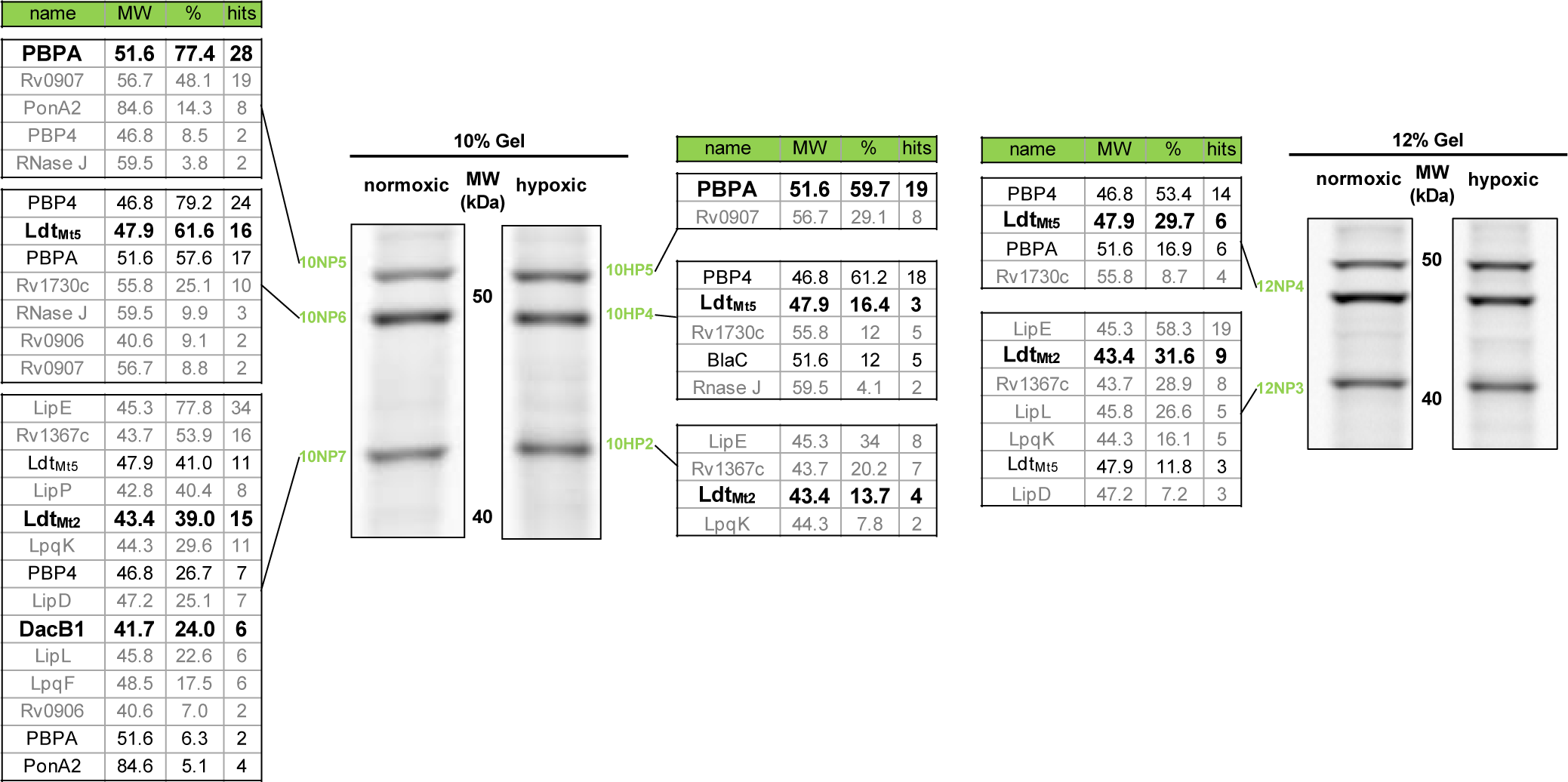
Identification of proteins associated with fluorescent bands observed between 40 and 50 kDa. Protein identifications for bands excised from the “middle region” of the 10% (left) and 12% (right) SDS PAGE gels. The most likely source of the fluorescent signal observed is provided in **Bold**. Known targets of β-lactams are indicated in black, while putative targets are in gray.

Interestingly, we observed a different banding pattern near 30 kDa for proteins obtained from normoxic versus hypoxic cultures (Figure 4B). We excised three of the 12% gel bands from hypoxic samples (10HS1, 10HS2, and 10HS4); all three contained BlaC peptides. Peptides from the carboxypeptidase DacB2 and other β-lactamases were also identified in these excised bands. Similar results were obtained from four bands excised from the 10% gel. This made the source of the signal ambiguous because it could be from any combination of BlaC, DacB2, or the putative β-lactamases. While it has been shown that BlaC represents the majority of β-lactamase activity under regular growth conditions^38^, similar studies have not been conducted with non-replicating bacteria. The presence of non-BlaC β-lactamases in dormant bacteria could alter their resistance profile toward various classes of β-lactam antibiotics.

The bands representing the lowest molecular weight proteins were excised from a 12% gel. Based on size, the signal near 15 kDa (band ID 12NP8) is most likely derived from Mero-Cy5 bound to Ldt_Mt3_ (28.7 kDa). Ldt_Mt3_ was also found in a band excised from another normoxic sample (12NS4). We sometimes observed other fluorescent bands in the 15-20 kDa region of the gel (see Figure 6), but those proteins were less stable to storage and electrophoresis and we were unable to identify them.

**Figure 6.**
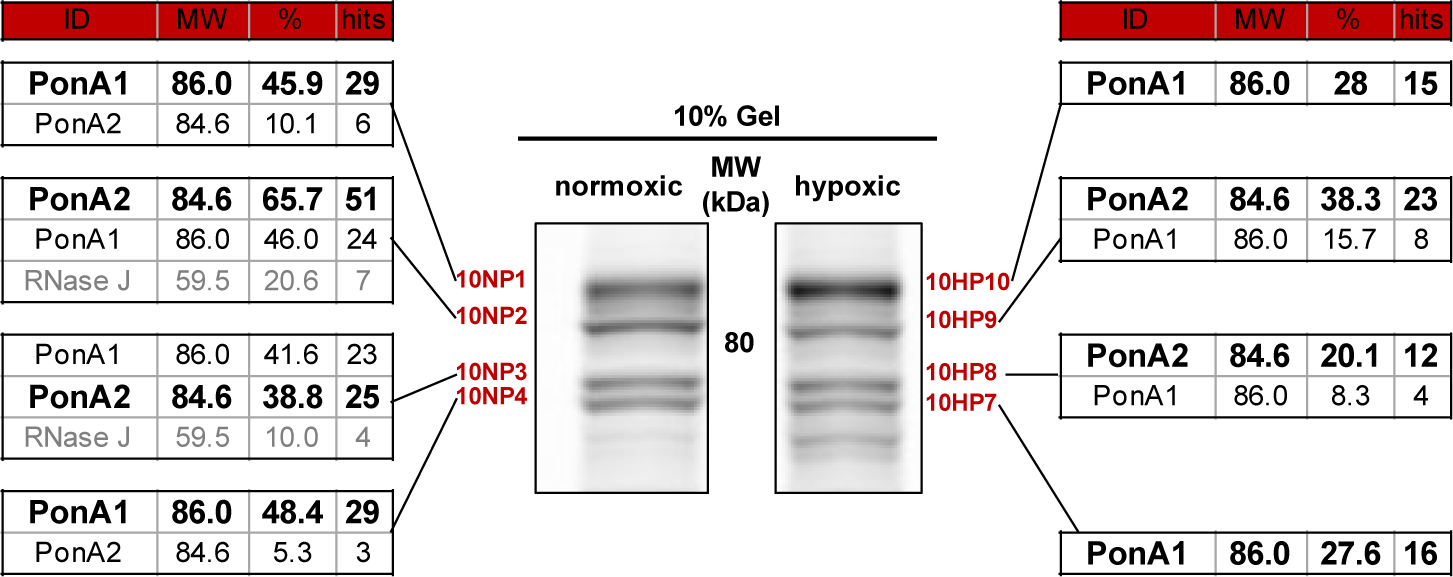
Identification of proteins associated with fluorescent bands near 80 kDa. Protein identifications for bands excised from a 10% SDS PAGE gel. Two PBPs, PonA1 and PonA2, were identified by LC-MS/MS. The most likely source of the fluorescent signal observed is provided in **Bold**.

Next we analyzed the middle region of the gel, which contained three fluorescent bands between 40 and 50 kDa (Figure 5). The two bright bands near 50 kDa were observed in both hypoxic and normoxic samples. These bands provided peptide hits from nine proteins. Based on molecular weight, we deduced that these two bands most likely contain PBPA (51.6 kDa), PBP4 (46.8 kDa), or LDT_Mt5_ (47.9 kDa). We believe that the upper band is PBPA based on % coverage and the number of peptide hits. When we compared the gel images from different β-lactam probes (see Figure 3), the band just below 50 kDa was only prominent with Mero-Cy5. This implies that the target can hydrolyze and release monobactam, cephalosporin, and/or penicillin based compounds—but not carbapenems^27,48^. We therefore believe that this signal was generated by Ldt_Mt5_.

Analysis of the band just above 40 kDa resulted in fourteen proteins with relevant Pfam notation. For these, eight were putative β-lactamases, four were PBPs (PBP4, DacB1, PBPA, and PonA2) and two were LDTs (Ldt_Mt5_ and Ldt_Mt2_). By molecular weight, DacB1 (41.7 kDa) and Ldt_Mt2_ (43.4 kDa) are the most likely to be present in this region. Ldt_Mt2_ was identified from three bands (10NP7, 10HP2, and 12NP3), corroborating its presence near 40 kDa.

From the “high region”, we analyzed bands from near 80 kDa (Figure 6). We obtained peptide hits from normoxic and hypoxic samples, with good sequence coverage for both PonA1 and PonA2. We believe that the signal in the top and bottom of the four bands is from PonA1. The appearance of PonA1 in two locations is unsurprising because it is regulated by phosphorylation, which can influence protein migration by SDS-PAGE*^(52, 60)^*. In hypoxia, the top band (10HP10) is more fluorescent than the bottom one (10HP7), which may indicate that PonA1 is differentially modified in dormancy. We believe that the two middle bands (10NP2 and 10NP3) contain predominantly PonA2, with the lower band resulting from cleavage of a 3 kDa signal sequence^*(61)*^. The signal peptide was only identified in the upper band (10NP2).

To summarize, we used LC-MS/MS to identify five PBPs (PonA1, PonA2, PBPA, DacB1, and DacB2), BlaC, and three LDTs (LDT_Mt2_, LDT_Mt3_, LDT_Mt5_). We also found evidence of many putative β-lactamases. Future studies might use purified proteins or genetic knock-out strains to more definitively assign activities identified here. Alternatively, samples could be treated with a biotinylated ABP and affinity-enriched before MS-based analysis. That approach is useful for enriching rare targets before LC-MS/MS analysis. Overall, our results demonstrate the utility of mero-Cy5 for identifying β-lactam drug targets in both normoxic and hypoxic *Mtb*.

### Patterns of protein activity from hypoxic and normoxic *Mtb*

Our MS analysis suggested that the activity of PBPs and LDTs changes in response to hypoxia. This fits prior observations of that *Mtb* increases the LDT-mediated cross-links in dormancy*^(11, 62, 63)^*. We investigated enzyme regulation further using ABP-labeled lysates from hypoxic and normoxic (actively-replicating). In Figure 7, the most plausible protein identifications are annotated based on our MS analysis.

**Figure 7.**
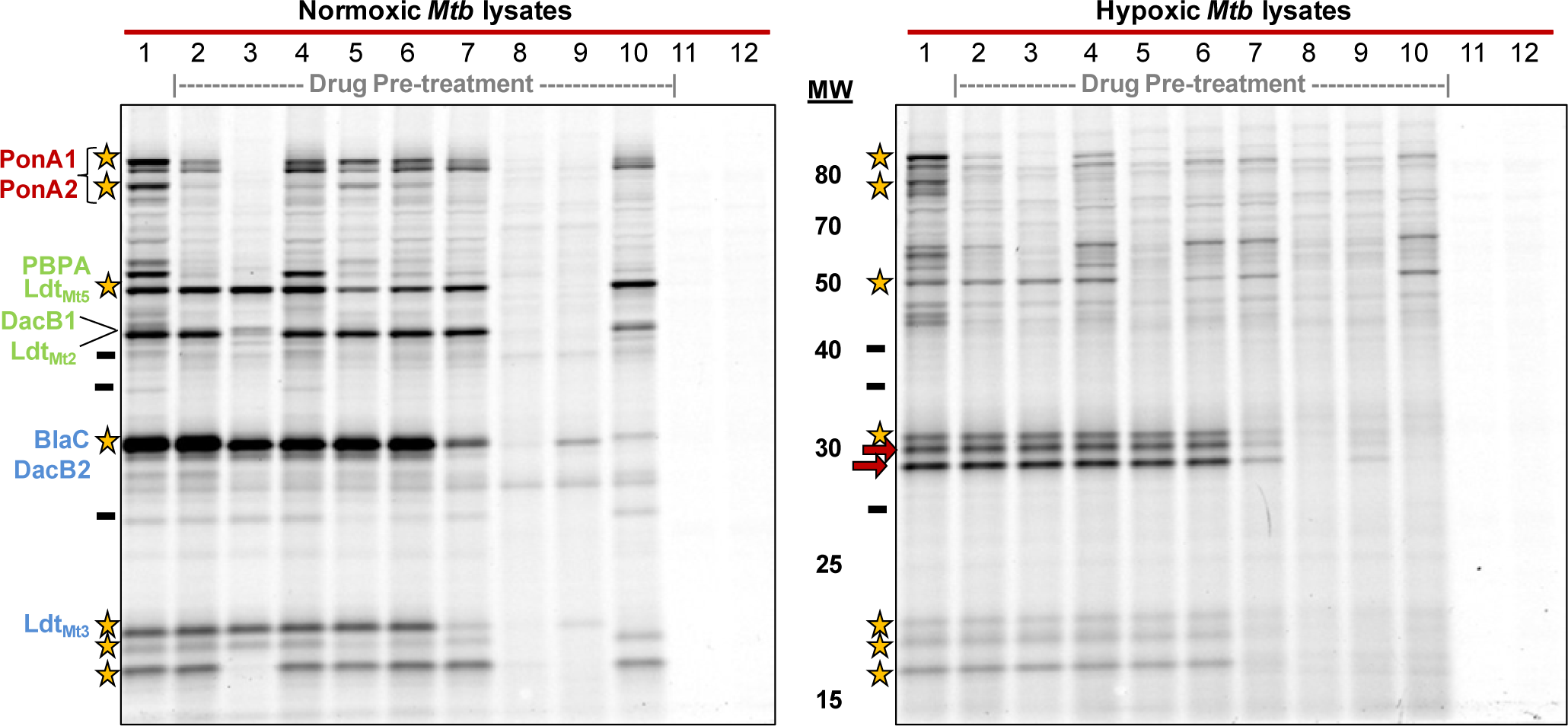
Mero-Cy5 reveals enzyme regulation and drug targets in lysates obtained from normoxic and hypoxic *Mtb*. Lane 1: No drug pre-treatment. Lanes 2-10: Drug pre-treatment. 2: Cephalexin; 3: Ceftriaxone; 4: Aztreonam; 5: Penicillin G; 6: Ampicillin; 7: Carbenicillin; 8: Meropenem; 9: Faropenem; 10: Clavulanate. Lane 11: Lysates *not* treated with Mero-Cy5. Lane 12: Lysates treated with a Cy5-(triazole)-butanoic acid. Putative protein identities were obtained by MS analysis. *Legend*: Gold stars denote enzymes that retain activity under hypoxic conditions. Red arrows indicate bands with enhanced activity in hypoxia. Black dash-marks indicate bands that are absent in hypoxia, but present under normoxia.

Mero-Cy5 revealed changes in PBP and LDT activity in dormancy. We found that many targets were present—if at lower intensity—in both growth conditions (stars;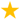). For example, PonA1, PonA2, LDT_Mt3_, and LDT_Mt5_ maintained their activity levels in dormancy. Others targets, particularly between 40 and 80 kDa, were less active in dormancy and at least three bands were absent (black dash-mark; **-**). The cephalexin probe indicated some up-regulation of faint bands in the 80 kDa region of the gel (**Figure S4**). These changes were also visible with Azt-Cy5, Mero-Cy5, and Bocillin-FL (**Figure S4**). It would be interesting to know if these changes are a result of post-translational modifications, such as phosphorylation^*(52)*^. Bocillin-FL probe offered little insight for targets below 50 kDa, as expected.

As noted in the previous section, we observed up-regulated enzyme activity near 30 kDa from hypoxic lysates. In particular, the ~28 kDa band was only distinct in hypoxic samples. This region was also notable in lysates treated with Azt-Cy5, which showed similar up-regulation of bands from 25-30 kDa under hypoxia (**Figure S4**). Our proteomics results indicate that DacB2 migrates in this region of the gel. This suggests that DacB2 could have enhanced activity under hypoxic conditions, an interesting result consistent with a prior proteomics study^*(40)*^. To summarize, our in-gel analysis provides evidence that the PBPs and LDTs are dynamically regulated in response to hypoxia.

### Drug susceptibility of PBPs and LDTs in *Mtb*

We set out to investigate antibiotic susceptibility in *Mtb* using ABPs. For these studies, we analyzed ABP-labeled lysates alongside lysates pre-treated with a selection of β-lactam antibiotics as inhibitors (Figure 6). Again, we studied lysates from dormant (hypoxic) and actively-replicating cultures. After drug-treatment, lysates were labeled with Mero-Cy5. As an additional control we exposed lysates to a sulfo-Cy5 conjugated to 5-hexynoic acid instead of a β-lactam (Lane 12). The absence of labeling with this compound indicated that the β-lactam, not Cy5, dictated target binding specificity. We tested cephalexin, ceftriaxone, aztreonam, penicillin G, ampicillin, carbenicillin, meropenem, faropenem, and clavulanate. All of these drugs except faropenem are prescribed for treating bacterial infections in the United States.

We were most interested in identifying bands that are targets of the carbapenems (i.e., meropenem and faropenem) because these drugs inhibit LDTs^*(22)*^. As expected, pre-treatment with meropenem (Lane 8), the closest structural analog to Mero-Cy5, eliminated most of the ABP’s labeling. We observed that meropenem completely inhibited a band that corresponds to Ldt_Mt5_, which is consistent with Lamichhane *et al.*’s finding that meropenem acylates the Ldt_Mt5_ active site^*(14)*^. Similar results were found with faropenem (Lane 9).

Ceftriaxone (Lane 3), a clinically-approved cephem, was more effective than cephalexin (Lane 2) at blocking Mero-Cy5 binding, although neither antibiotic was an effective inhibitor of BlaC, Ldt_Mt5_, or Ldt_Mt3_. The penams, penicillin G (Lane 5), ampicillin (Lane 6), and carbenicillin (Lane 7), were less effective at inhibiting targets than other drugs, but reduced Mero-Cy5 labeling of PBPA. BlaC, which retained some activity in dormancy, was inhibited by several drugs (Lanes 7-10), including clavulanate (Lane 10) and meropenem (Lane 8).

Antibiotic susceptibility was also analyzed using Azt-Cy5 and CephX-Cy5, and we observed similar inhibition profiles (**Figure S5**). Overall, our findings indicate that PBP and LDT activities can be reduced or eliminated in vitro using clinically-approved β-lactams.

## Conclusion

In conclusion, we used fluorescent β-lactam probes to identify active PBPs, LDTs, and β-lactamases in *Mtb*. In a direct comparison with Bocillin-FL, we found that Azt-Cy5, CephX-Cy5, and Mero-Cy5 enabled more targets to be detected in PAGE-resolved lysates. Although we collected data with Cy5-modified probes, we reported the synthesis of the alkyne derivatives because they enable facile modification with an assortment of reporters. For example, a biotinylated probe would enable enrichment of β-lactam targets before proteomic analysis^*(55)*^.

We used Mero-Cy5, a new probe, to identify 32 proteins, including transpeptidases (PonA1, PonA2, PBPA, LDT_Mt2_, LDT_Mt3_, LDT_Mt5_), carboxypeptidases (DacB1, DacB2, PBP4), BlaC, and numerous putative β-lactamases (see **Table S2**). We did not identify LDT_Mt1_ or LDT_Mt4_, although there is structural evidence that both LDTs bind carbapenems^*(15–17)*^. We found evidence that post-translational modifications alter the migration patterns observed in-gel. In future work, it would be interesting to study in more detail how various post-translational modifications alter PBP^*(52)*^ and LDT activities during different stages of infection.

We used Mero-Cy5 to investigate enzyme regulation in dormant and actively-replicating samples. We found numerous changes in fluorescent banding patterns and intensity between the two states. For example, we found that some bands (25-30 kDa) were up-regulated under hypoxia. A priority for future work is to identify the exact source of that activity because it is unclear if those bands are attributable to BlaC. Lastly, we treated lysates with a set of β-lactam drugs in clinical use today and found that meropenem and faropenem were both good inhibitors of BlaC, PBPs, and LDTs. Overall, we demonstrated that mero-Cy5 is a useful probe for target identification, analysis of enzyme regulation, and determining enzyme inhibition by various β-lactam drugs.

## Methods

### Synthesis of activity-based probes

Bocillin FL was obtained from ThermoFisher Sci. Azt-alkyne, CephX-alkyne, and Mero-alkyne were synthesized from the parent β-lactam, as described in the Supporting Information. The fluorescent probes were generated in vitro via Cu-catalyzed click reaction (1h, room temperature) of the alkyne (100 µM) with sulfo-Cyanine5-azide (100 µM), 1.25 mM THTPA/0.25 mM CuS0_4_, and sodium ascorbate (15 mM) in HEPES buffer (pH 7.3).

### Mtb culture conditions

*Mtb* mc^2^6020 was grown in Middlebrook 7H9-OADC medium supplemented with lysine, pantothenate, and casamino acids. For normoxic cultures the flasks were maintained at 37 °C with 100 rpm shaking under atmospheric conditions. For hypoxic cultures, mid-log phase cultures were diluted to OD_600_ 0.4 with additional medium and grown as standing cultures under 1% O_2_ and 5% CO_2_ at 37 °C.

### Preparation of lysates

*Mtb* grown under normoxic conditions was harvested at mid-log phase (OD_600_ = 0.9–1.2). *Mtb* grown under hypoxic conditions was harvested at fixed time-points (OD_600_ = 0.4–0.8). All cells were collected by centrifugation (10 min, 4,000 x g, 4 °C) and washed twice with PBS containing 0.05% Tween 80. Pellets were resuspended in detergent-free lysis buffer [50 mM Tris (pH 7.5 at 4 °C), 50 mM NaCl, 0.5 mM CaCl_2_, 0.5 mM MgCl_2_], lysed by mechanical disruption, and pelleted by centrifugation (10 min, 4,000 x g, 4 °C). The supernatant was transferred to a separate tube and the beads and cell debris resuspended in an equivalent volume of detergent containing lysis buffer [50 mM Tris (pH 7.5 at 4 °C), 50 mM NaCl, 0.5 mM CaCl_2_, 0.5 mM MgCl_2_, 0.4% triton X-100] and the mechanical disruption and centrifugation steps were repeated. The combined supernatants were pelleted by centrifugation (10 min, 4,000 x g, 4 °C) to remove insoluble debris and sterilized by filtration (0.2 μm PES membrane).

### Lysate labeling and imaging

Lysates were diluted with detergent containing lysis buffer [50 mM Tris (pH 7.5 at 4 °C), 50 mM NaCl, 0.5 mM CaCl_2_, 0.5 mM MgCl_2_, 0.2% triton X-100] to normalize the total protein concentration. For samples pre-treated with antibiotics, the antibiotic was added for 15 min prior to probe-labeling. Lysates were labeled with the Cy5-modified probe (5 μM) for 1 h at rt. Lysates were resolved by SDS-PAGE, washed, fixed, and imaged on a Typhoon multi-mode imager.

### Band excision and LC-MS/MS proteomics

All bands were excised from an SDS-PAGE gel based on the fluorescent image and stored frozen. Gel slices were processed to remove SDS, reduced, and methylated. Proteins were trypsin-digested and the resulting peptides were concentrated. LC-MS/MS based proteomics was performed by the OHSU Proteomics Shared Resource Facility.

## Supporting information

The Supporting Information includes detailed methods and Figures S1-S7. An Excel file containing Tables S1 and S2 are available by request.

## Acknowledgments

Funding for this research was provided by the Knight Cancer Institute and the OHSU School of Medicine. S.R.L. was supported by an NIH T32 training grant (T32-AI07472). We are grateful to Dr. Gyanu Lamichhane (JHU) and Dr. Clifton Barry (NIH) for helpful discussions, Scotland Farley (OHSU) for synthesizing the aztreonam probe, and Dr. Kyle Gee (ThermoFisher Sci) for providing Bocillin FL.

