## Supplementary Information for "Investigating β-lactam drug targets in *Mycobacterium tuberculosis* using chemical probes"

### Supplementary Methods

Unless otherwise stated, reactions were magnetically stirred in flame-dried glassware under an atmosphere of nitrogen. Anhydrous solvents were purchased in septum-sealed bottles and stored under nitrogen. Semi-dry DMF was stored over 4 Å molecule sieves. $Et_3N$ was stored over $K_2CO_3$ in a desiccator. All other chemicals were purchased from commercial sources and used as received. Reactions were monitored by thin-layer chromatography (TLC) on glass-backed silica gel plates (Silicycle 60 Å, 250 μM). Reverse phase column chromatography was performed with the indicated solvents on a Biotage Isolera One automated chromatography system fitted with a 12 g KP-C18-HS column.

Mass spectra were acquired at Portland State University's bioanalytical mass-spectrometry facility on a ThermoElectron LTQ-Orbitrap Discovery high-resolution mass spectrometer with electrospray ionization (ESI).

NMR spectra were taken at ambient temperature at Portland State University's NMR facility. $^1$H-NMR data were obtained in the specified solvent on a Bruker Avance II at 400 MHz. $^{13}$C-NMR data were obtained on the same instrument in the specified solvent at 101 MHz. Spectra were calibrated to the residual solvent peak. Chemical shifts are reported in ppm. Coupling constants (*J*) are reported in Hertz (Hz) and rounded to the nearest 0.1 Hz. Multiplicities are defined as: s = singlet, d = doublet, dd = doublet of doublets, t = triplet, td = triplet of doublets, quin = quintet, m = multiplet, br s = broad singlet.

#### Synthesis of (*Z*)-3-(2-(2-aminothiazol-4-yl)-2-((2,2-dimethyl-3-oxo-3-(prop-2-yn-1-ylamino)propyl)imino)acetamido)-2-methyl-4-oxoazetidine-1-sulfonate (aztreonam-alkyne)

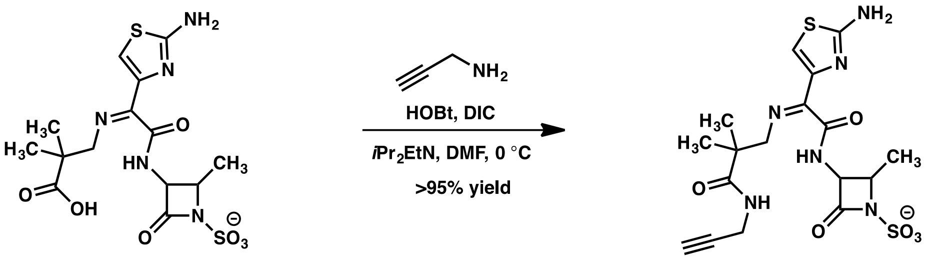


**aztreonam** **aztreonam-alkyne**

In a 10 mL round-bottom flask with two necks, aztreonam (0.0310 g, 0.07 mmol, 1 equiv) was dissolved in anhydrous DMF (2 mL, 0.035 M). The flask was cooled to 0 °C and HOBt (0.0153 g, 0.10 mmol, 1.4 equiv), DIC (16 μL, 0.102 mmol, 1.5 equiv), and N,N-diisopropylethylamine (30 μL, 0.17 mmol, 2.5 equiv) were added. After stirring for 30 min, propargylamine amine (6.6 μL, 0.103 mmol, 1.5 equiv) was added. The reaction mixture was allowed to gradually warm to room temperature (rt) and stirring continued for 3 d. The reaction mixture was concentrated *in vacuo* and purified by column chromatography on $SiO_2$ (85% EtOAc/15% 1:1 $H_2O$/AcOH → 65% EtOAc/35% 1:1 $H_2O$/AcOH). The product was an orange solid (0.0322 g, 96% yield) whose spectral data matched the literature values[1].

### Generation of 5-hexynoic acid chloride

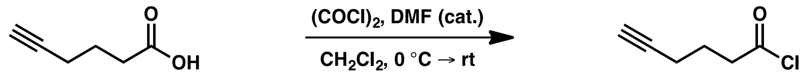


In a 10 mL pear-shaped flask, a solution of 5-hexynoic acid (0.25 mL, 2.27 mmol, 1 equiv) in $CH_2Cl_2$ (1.5 mL) was cooled to 0 °C. Oxalyl chloride (0.45 mL, 5.25 mmol, 2.3 equiv) was added, followed by a catalytic drop of dry DMF. After stirring for 20 min at 0 °C the flask was allowed to warm and stirred at rt for 40 min. The resulting yellow solution was concentrated *in vacuo* to a red-orange oil. The acid chloride was diluted with dry DMF and the solution kept under nitrogen.

### Synthesis of (6R,7R)-7-((R)-2-(hex-5-ynamido)-2-phenylacetamido)-3-methyl-8-oxo-5-thia-1-azabicyclo[4.2.0]oct-2-ene-2-carboxylic acid (cephalexin-alkyne)

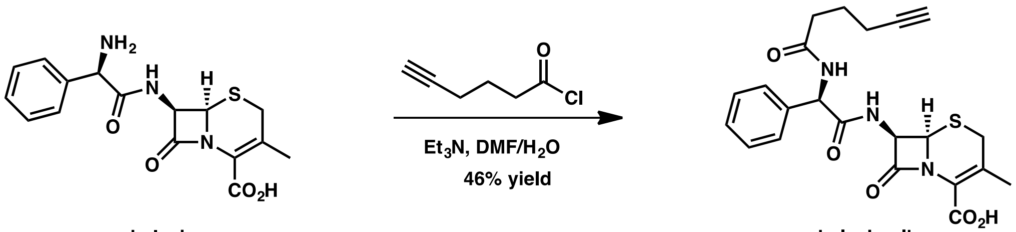


cephalexin — cephalexin-alkyne

Cephalexin monohydrate (0.0842 g, 0.23 mmol, 1 equiv) was suspended in semi-dry DMF (1 mL) in a 2-dram vial. A solution of 5-hexynoic acid chloride (~1 M, 0.2 mL, 0.20 mmol, 0.87 equiv) was added to this vial, followed by $Et_3N$ (0.15 mL, 1.08 mmol, 4.7 equiv), then $H_2O$ (0.15 mL). The vial was capped and the reaction mixture stirred at rt. Additional hexynoic acid chloride solution was added at 40 and 110 min (2x 0.1 mL, 0.1 mmol, 0.43 equiv). Additional $Et_3N$ (0.05 mL, 0.36 mmol, 1.6 equiv) was added at 2.5 h and $H_2O$ (0.25 mL) at 3 h. After 4 h the reaction mixture was cooled to -80 °C for overnight storage. After warming to rt, additional $H_2O$ was added and the reaction mixture purified via reverse phase column chromatography (0 → 100% MeCN in $H_2O$, with 0.1% TFA). Concentration *in vacuo* gave an orange solid identified as cephalexin-N-alkyne (0.0488 g, 46% yield).

$^1$H-NMR (400 MHz; DMSO-d$_6$): δ 13.23 (br s, 1H), 9.25 (d, *J* = 8.2 Hz, 1H), 8.56 (d, *J* = 8.2 Hz, 1H), 7.45-7.42 (m, 2H), 7.34-7.25 (m, 3H), 5.67 (d, *J* = 8.3 Hz, 1H), 5.62 (dd, *J* = 8.2, 4.7 Hz, 1H), 4.96 (d, *J* = 4.7 Hz, 1H), 3.46 (d, *J* = 19.1 Hz, 2H), 3.28 (d, *J* = 18.1 Hz, 1H), 2.77 (t, *J* = 2.6 Hz, 1H), 2.31 (t, *J* = 7.2 Hz, 2H), 2.14 (td, *J* = 7.3, 2.6 Hz, 2H), 1.98 (s, 3H), 1.66 (quin, *J* = 7.3 Hz, 2H).

### Synthesis of (4*R*,6*S*)-3-(((3*R*,5*R*)-5-(dimethylcarbamoyl)-1-(hex-5-ynoyl)pyrrolidin-3-yl)thio)-6-((*R*)-1-hydroxyethyl)-4-methyl-7-oxo-1-azabicyclo[3.2.0]hept-2-ene-2-carboxylic acid (meropenem-alkyne)

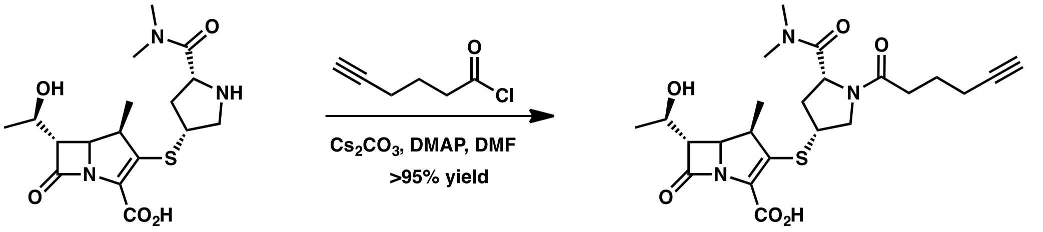


meropenem — meropenem-alkyne

Meropenem trihydrate (0.0441 g, 0.10 mmol, 1 equiv) was dissolved in semi-dry DMF (0.5 mL) in a 1 dram vial. $Cs_2CO_3$ (0.1016 g, 0.31 mmol, 3.1 equiv) and a catalytic chip of DMAP were added and the solution stirred at rt. A solution of 5-hexynoic acid chloride (~2 M, 0.1 mL, 0.2 mmol, 2 equiv) was added drop-wise and the resulting heterogeneous mixture stirred at rt. After 30 min, additional 5-hexynoic acid chloride solution was added (~2 M, 0.05 mL, 0.1 mmol, 1 equiv). After 45 min additional $Cs_2CO_3$ (0.0501 g, 0.15 mmol, 1.5 equiv) was added. At 1 h, the reaction mixture was transferred to a microcentrifuge tube and the Cs2$CO_3$ pelleted. The supernatant was removed and triturated in batches with 0.5 mL $Et_2O$ for every 0.1 mL of supernatant. After centrifugation the pellets (a brown tar) were combined, washed with $Et_2O$ (2 x 1 mL), and dried briefly under vacuum. Once the brown tar was a solid mass it was crushed and left under vacuum to give an orange-tan solid (0.0569 g, 118% yield), which retained some DMF and/or $H_2O$.

LC-MS m/z calculated for [M+H$^+$]: 478.2006 [$C_{23}H_{32}N_3O_6S$], found: 478.2098

#### Cu-catalyzed azide-alkyne click (CuAAC) reaction

For all CuAAC reactions, the THPTA and $CuSO_4$ were combined in advance in $H_2O$. This 10x stock solution contained 12.5 mM THPTA and 2.5 mM $CuSO_4$. It was kept at –20 °C for long-term storage.

The following compounds were combined in a microcentrifuge tube containing 7 µL 10 mM HEPES buffer (pH 7.3) (in order of addition): β-lactam alkyne probe (4 µL, 500 µM in HEPES, final concentration 100 µM), Sulfo-Cyanine5-azide (Lumiprobe; 5 µL, 400 µM in HEPES, final concentration 100 µM), THPTA ligand / $CuSO_4$ mixture (2 µL, 12.5 mM THPTA / 2.5 mM $CuSO_4$ in $H_2O$, final concentration 1.25 mM THPTA / 250 µM $CuSO_4$), and sodium ascorbate (2 µL, 150 mM in $H_2O$, final concentration 15 mM). The reaction was incubated for 1 h at room temperature in the dark. The final reaction mixture was assumed to be 100 µM in β-lactam-Cy5. If not used immediately the sample was stored frozen at –20 °C. *Note:* for the Cy5 probe control, the β-lactam alkyne was replaced with 5-hexynoic acid.

#### Mycobacterial culture conditions

All species were handled as BSL-2 pathogens.

Middlebrook 7H9 broth (271310) and Bacto CAS amino acids (223120) were purchased from Becton Dickinson. OADC and ADC were made during media preparation from the following components: oleic acid (S25451A), catalase (S25239A), and sodium chloride (BP358) purchased from Fisher Scientific. BSA fraction V (fatty acid free) was purchased from Research Products International (A30075-100.0). Dextrose was purchased from Sigma-Aldrich (D9434). All liquid media was sterile filtered (0.2 µm PES) and stored at 4 °C.

Cells were thawed from frozen stocks (stored at –80 °C in 6% glycerol). *Mtb* mc$^2$6020 (ΔlysA ΔpanCD mutant[2]) was cultured in 7H9-OADC medium (7H9 broth, 0.5% glycerol, 0.05% Tween 80, 10% OADC) supplemented with 80 µg/mL lysine, 24 µg/mL pantothenate, and 0.2% casamino acids. Cultures were grown at 37 °C with 100 rpm shaking. *Mtb* grown under normoxic conditions was harvested at $OD_{600}$ = 0.9–1.2. Cells were collected by centrifugation (10 min, 4,000 x g, 4 °C). Pellets were washed twice with PBS containing 0.05% Tween 80 in order to remove BSA. Pellets were stored at –30 °C.

*M. smegmatis* mc$^2$155 (ATCC 700084) was cultured in 7H9-ADC medium (7H9 broth, 0.5% glycerol, 0.05% Tween 80, 10% ADC) at 37 °C with 100 rpm shaking. *M. marinum* Aronson (ATCC 927) was grown in the same medium at 30 °C with 100 rpm shaking in the dark. *M. smegmatis* was harvested at $OD_{600}$ = 1.3. *M. marinum* was harvested at $OD_{600}$ = 0.7. Cells were collected by centrifugation (10 min, 4,000 x g, 4 °C), washed twice with PBS containing 0.05% Tween 80, and the pellets were stored at –30 °C.

#### Hypoxia induced dormancy of *Mtb* mc$^2$6020

Cultures were grown to an $OD_{600}$ between 0.5 and 1.0 and then diluted to an $OD_{600}$ of 0.4 in 7H9 medium (see above). Cultures (50–150 mL) were grown at 37 °C in standing flasks with vented caps in a Heracell 150i incubator (Thermo Scientific) at 1% $O_2$ and 5% $CO_2$. Depending on the volume of the culture, methylene blue (1.5 µg/mL) de-colorization in control cultures indicated oxygen depletion between 14 and 28 days, as previously reported[3]. Cultures were harvested by centrifugation (10 min, 4,000 x g, 4 °C) at 33 days (methylene blue de-colored at 28 days, OD = 0.8) for all experiments excepting the antibiotic panels (Figure 7; Figure S5), which were harvested at 15 days (OD = 0.4). The pellets were washed twice with PBS containing 0.05% Tween 80 in order to remove BSA, and stored at –30 °C.

**Preparation of lysates**

Pellets were thawed and resuspended in detergent-free lysis buffer [50 mM Tris (pH 7.5 at 4 °C), 50 mM NaCl, 0.5 mM $CaCl_2$, 0.5 mM $MgCl_2$] and lysed by mechanical disruption on a BioSpec Mini Beadbeater (6 x 45 s pulses with cooling on ice between rounds) using 0.1 mm zirconia/silica beads (BioSpec Products). The samples were then pelleted by centrifugation (10 min, 4,000 x g, 4 °C). The supernatant was transferred to a separate tube and the beads and cell debris resuspended in an equivalent volume of detergent containing lysis buffer [50 mM Tris (pH 7.5 at 4 °C), 50 mM NaCl, 0.5 mM $CaCl_2$, 0.5 mM $MgCl_2$, 0.4% triton X-100] and the mechanical disruption and centrifugation steps were repeated. The combined supernatants were pelleted by centrifugation (10 min, 4,000 x g, 4 °C) to remove insoluble debris and zirconia/silica beads. Lysates were sterilized by filtration through a 0.2 µm PES membrane (Agela Technologies 25 mm syringe filter with glass fiber pre-filter, AS052520-G).

Total protein quantification was determined by BCA assay (Pierce, 23225). If the lysates were not used immediately, 6% glycerol was added (10% additional volume using a 60% glycerol solution) and the samples stored at –80 °C.

**Analysis of labeled lysates by SDS-PAGE**

Lysates or enriched samples were thawed on ice and diluted with detergent containing lysis buffer [50 mM Tris (pH 7.5 at 4 °C), 50 mM NaCl, 0.5 mM $CaCl_2$, 0.5 mM $MgCl_2$, 0.2% triton X-100] to give a normalized protein concentration across samples for a particular experiment. For samples pre-treated with antibiotics, the antibiotic (as a 5−10x stock in HEPES) was added and the reaction incubated for 15 min at room temperature in the dark. For all samples, the probe (clicked to sulfo-Cyanine5-azide, Lumiprobe) was added to a final concentration of 5 µM and the sample incubated for 1 h at rt in the dark. Reactions were quenched by the addition of 5x SDS-PAGE loading dye containing TCEP. Samples were heated to 95 °C for 5−10 min and 5 µg of total protein from each sample was separated on a Bio-Rad Criterion Bis-Tris gel with XT-MOPS running buffer. Gels were washed 3x briefly with $H_2O$ and fixed in 20% MeOH/10% AcOH/70% $H_2O$ to remove unreacted probe. After re-equilibrating in $H_2O$, fluorescent images were taken with a Typhoon 9400 variable mode imager (Amersham Biosciences). Adjustment of image brightness/contrast was performed with imageJ.

**Band excision for LC-MS/MS-based proteomics**

Lysates were fractionated before analysis. First, lysates were centrifuged overnight at 21,130 x g (4 °C). The resulting pellets were enriched in both cell-wall and membrane components[4]. Pellets were resuspended by vigorous pipetting in detergent containing lysis buffer [50 mM Tris (pH 7.5 at 4 °C), 50 mM NaCl, 0.5 mM $CaCl_2$, 0.5 mM $MgCl_2$, 0.2% triton X-100]. Total protein quantification for both the resuspended pellet and the supernatant was determined by BCA assay (Pierce, 23225).

Samples were prepared as described in the "Analysis of labeled lysates by SDS-PAGE" section up to the additional of loading dye. After heating at 60 °C for 20 min the samples were stored at –30 °C. Before running a gel, an additional aliquot of 5x SDS-PAGE loading dye containing TCEP was added and the samples heated to 95 °C for 5−10 min. For the lowest molecular weight bands, a 12% Bis-Tris gel was used with 20 µg of total protein loaded in the lanes to be excised. On the 12% gel, 5 µg of total protein was used for adjacent lanes to enable more precise band visualization. A 10% gel was used for the higher MW bands excised. Three adjacent lanes in a 10% gel were used per sample: two contained 10 µg of total protein per lane and one contained 5 µg (for better band visualization); all three lanes were excised together. After electrophoresis, the gels were washed briefly 3x with $H_2O$ and fixed in 20% MeOH/10% AcOH/70% $H_2O$ to remove unreacted probe. After re-equilibrating in $H_2O$, fluorescent images were taken with a Typhoon 9400 variable mode imager (Amersham Biosciences).

Bands were excised using a clean razor blade and stored in microcentrifuge tubes at –30 °C before processing.

#### Band Processing I

**Samples:** 10HS1, 10HS2, 10HS4, 10NP1, 10NP2, 10NP3, 10NP4, 10NP5, 10NP6, 10NP7, 10NP8, 12NP8, 12NP9, 12NS4, 12aztHS1, 12aztHS2, 12aztHS3, 12cephHS, 12cephNS

After thawing, the gel slices were broken up into pieces in their tubes. The gel pieces were then washed 3x with 100 µL of AMBIC [50 mM ammonium bicarbonate buffer, pH 8 (freshly prepared)]. The gel pieces were dehydrated by washing 3−4x with 100 µL acetonitrile. The dehydrated gel pieces were stored at 4 °C overnight. The gel pieces were rehydrated in 150 µL of 10 mM DTT (in AMBIC) for 30 min at 56 °C. The gel pieces were dehydrated by washing 3x with 100 µL acetonitrile. The gel pieces were rehydrated in 150 µL of 55 mM iodoacetamide (in AMBIC) for 30 min in the dark at room temperature. The gel pieces were washed 2x with 200 µL of AMBIC. The gel pieces were dehydrated by washing 1x 200 µL and 3x 100 µL with acetonitrile.

Sequencing grade trypsin (Promega V5111A) was resuspended in trypsin resuspension buffer (Promega V542A) at 20 µg/µL. Next, 2 µL of this solution was diluted into 998 µL of digestion buffer (0.01% freshly prepared ProteaseMAX in AMBIC, Promega V2071) to give a 2 ng/µL solution of trypsin. For each sample, 50 µL (100 ng of trypsin) of this solution was added and samples were incubated at rt for 10 min. An additional 50 µL of digestion buffer was added and the samples incubated at 50° C for 1 h. Digestion was halted with 5 µL formic acid (Fisher A117-50).

The samples were pelleted by centrifugation (17,000 x g, 5 min) and the supernatants collected in microcentrifuge tubes (Sarstedt 72.692.005). The gel pieces were then combined with 50 µL of 60% acetonitrile (Fisher Scientific A998-4) and 0.1% trifluoroacetic acid (Fisher Scientific O4902-100) in $H_2O$ and sonicated in a water bath for 15 min. The samples were pelleted by centrifugation (17,000 x g, 10 min) and the supernatants collected and stored at −30 °C prior to analysis.

Concentration by SpeedVac was performed by the OHSU Proteomics Shared Resource Facility.

#### LC-MS/MS-based proteomics for excised bands I

Each prepared/digested sample was then dissolved in 20 µL of 5% formic acid and analyzed using an Orbitrap Fusion mass spectrometer configured with an EasySpray NanoSource (Thermo Scientific). Digests were loaded onto an Acclaim PepMap 0.1 x 20 mm NanoViper C18 peptide trap (Thermo Scientific) for 5 min at a 5 µL/min flow rate in a 2% acetonitrile, 0.1% formic acid mobile phase and peptides separated using a PepMap RSLC C18, 2 µm particle, 75 µm x 25 cm EasySpray column (Thermo Scientific) using a 7.5–30% acetonitrile gradient over 60 min in mobile phase containing 0.1% formic acid and a 300 nL/min flow rate using a Dionex NCS-3500RS UltiMate RSLC nano UPLC system. Survey scans from 400-1600 m/z were performed in the Orbitrap mass analyzer at 120,000 resolution, AGC target of $4 \times 10^5$, maximum injection time of 50 ms, and a m/z= 445.12 polysiloxane ion lock mass.

Data-dependent MS2 scans were performed in the linear ion trap using HCD following isolation with the instrument's quadrupole. Data dependent MS/MS scans used a cycle time of 3 sec between survey scans, a mono isotopic peak determination filter with a peptide setting, intensity threshold of $5 \times 10^3$, dynamic exclusion of 30 sec with a +/- 10 ppm mass tolerance, charge state filter to ignore +1 ions, quadrupole isolation window of 1.6 m/z, HCD collision energy of 30%, rapid scan rate, AGC target of $1 \times 10^4$, and maximum injection time of 35 ms.

Comet (version 2016.01, revision 1)[5] was used to search MS2 spectra against *Mycobacterium tuberculosis* (strain H37Rv) sequences from UniProt (downloaded: June 2019). The database also contained concatenated sequence-reversed entries to estimate error thresholds and 179 common contaminant sequences and their reversed forms as well as the additional sequence for the *Mtb* protein CrfA (see below). The database processing was performed with python scripts available at https://github.com/pwilmart/fasta_utilities.git and Comet results processing used the PAW pipeline[6] from https://github.com/pwilmart/PAW_pipeline.git.

Comet searches for all samples were performed with trypsin enzyme specificity. Average parent ion mass tolerance was 1.25 Da to improve estimates of peptide false discovery. Monoisotopic fragment ion mass tolerance was 1.0005 Da. A static modification of +57.02146 Da was added to all cysteine residues. A variable modification of +15.9949 Da on methionine residues was also allowed.

Scaffold (version Scaffold_4.8.4, Proteome Software Inc., Portland, OR) was used to validate MS/MS based peptide and protein identifications. Peptide identifications were accepted if they could be established at greater than 95.0% probability (peptide FDR = 0.6%) by the Peptide Prophet algorithm[7] with Scaffold delta-mass correction. Protein identifications were accepted if they could be established at greater than 99.0% probability (protein FDR = 0.0%) and contained at least 2 identified peptides. Protein probabilities were assigned by the Protein Prophet algorithm[7]. Proteins that contained similar peptides and could not be differentiated based on MS/MS analysis alone were grouped to satisfy the principles of parsimony.

The CrfA sequence used was[8]:
VLGGGDLSTLLPGLPAGHEYHFVEIEQVGHLAGGNQVAVVDGVERPTHDSQSTPMHDWPAYWSILPQF
RSRQLPASLSEQPLIRQHDWLFLSLCVIILVSFRVRSRSSPTGRDWVSRPSPVTPSASGMAYR

#### Band Processing II

**Samples:** 10HP1, 10HP2, 10HP4, 10HP5, 10HP7, 10HP8, 10HP9, 10HP10, 10HS3, 12HS1, 12HS2, 12HS3, 12NP3, 12NP4

These samples were prepared by the OHSU Proteomics Shared Resource Facility according to the following protocol. Gel bands were digested using the "In-Gel Digestion Protocol for Low Protein Amounts" procedure described in section 5 of the Promega ProteaseMax Surfactant, Trypsin Enhancer Technical Manual.

#### LC-MS/MS-based proteomics for excised bands II

The peptides were separated using the same nano UPLC conditions as above using a 60 min gradient, except a Symmetry C18 trap and 75 μm x 250  BEH 130 C18 column with 1.7 μm particles (Waters) was used and peptides analyzed using a Q-Exactive HF mass spectrometer and Nano Flex Ion Spray Source (Thermo Scientific) fitted with a 20 μm stainless steel nano-bore emitter spray tip. Survey mass spectra were acquired in m/z 375−1400 at 120,000 resolution, AGC target of 3 x $10^6$,maximum injection time of 50 ms, and a m/z= 445.12 polysiloxane ion lock mass. Data-dependent MS/MS acquisition selected the top 10 most abundant precursor ions using an isolation width of 1.2 m/z, normalized collisional energy of 30, AGC setting of 1 x $10^5$, maximum ion time of 100 ms, minimum AGC target of 5 x $10^3$, exclusion of +1 ions, and dynamic exclusion in auto mode.

MS/MS spectra were searched as above, with the only change in parameters being the peptide FDR = 0.4%.

**Supplementary data.**

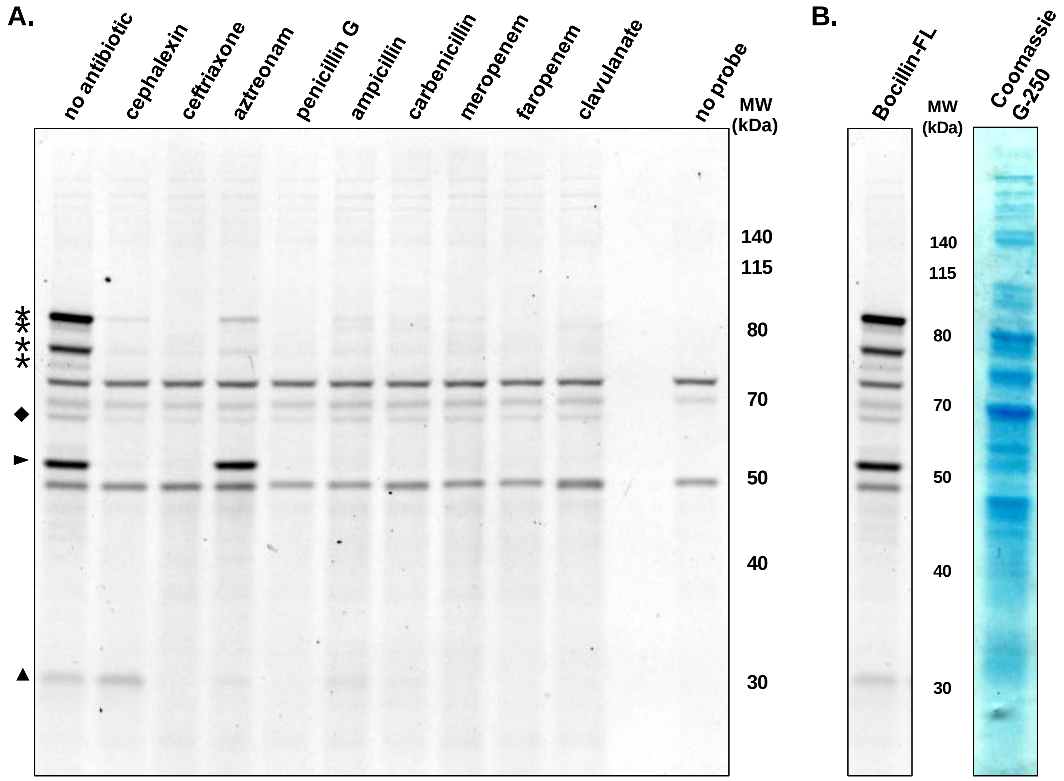


**Figure S1. Bocillin-FL selectively labels proteins in *Mtb* lysates. A.** Whole cell lysates from *Mtb* mc[2]6020 were treated with the indicated antibiotic (500 μM) for 15 min at room temperature. Bocillin-FL (5 μM) was added to all samples except the "no probe" control. The reactions were incubated for 1 h. Reactions were quenched by the addition of loading dye, boiled, and separated by SDS-PAGE (12% Bis-tris gel with XT-MOPS running buffer). The gel was washed and fixed overnight (20% MeOH/10% AcOH/70% $H_2O$) to remove excess probe. The gel image was taken at $\lambda_{ex}/\lambda_{em}$ = 488/520BP40 (nm) on a Typhoon Multimode Imager. ★ indicates bands that were inhibited by all tested compounds. ◆ indicates a band that was resistant to inhibition by the tested compounds. ► indicates a band that was significantly inhibited by all compounds except aztreonam. ▲ indicates a band with partial or complete inhibition with all compounds except cephalexin. The “no probe” control shows evidence of three autofluorescent proteins. **B.** Bocillin-FL labeling does not correlate with coomassie staining for total protein.

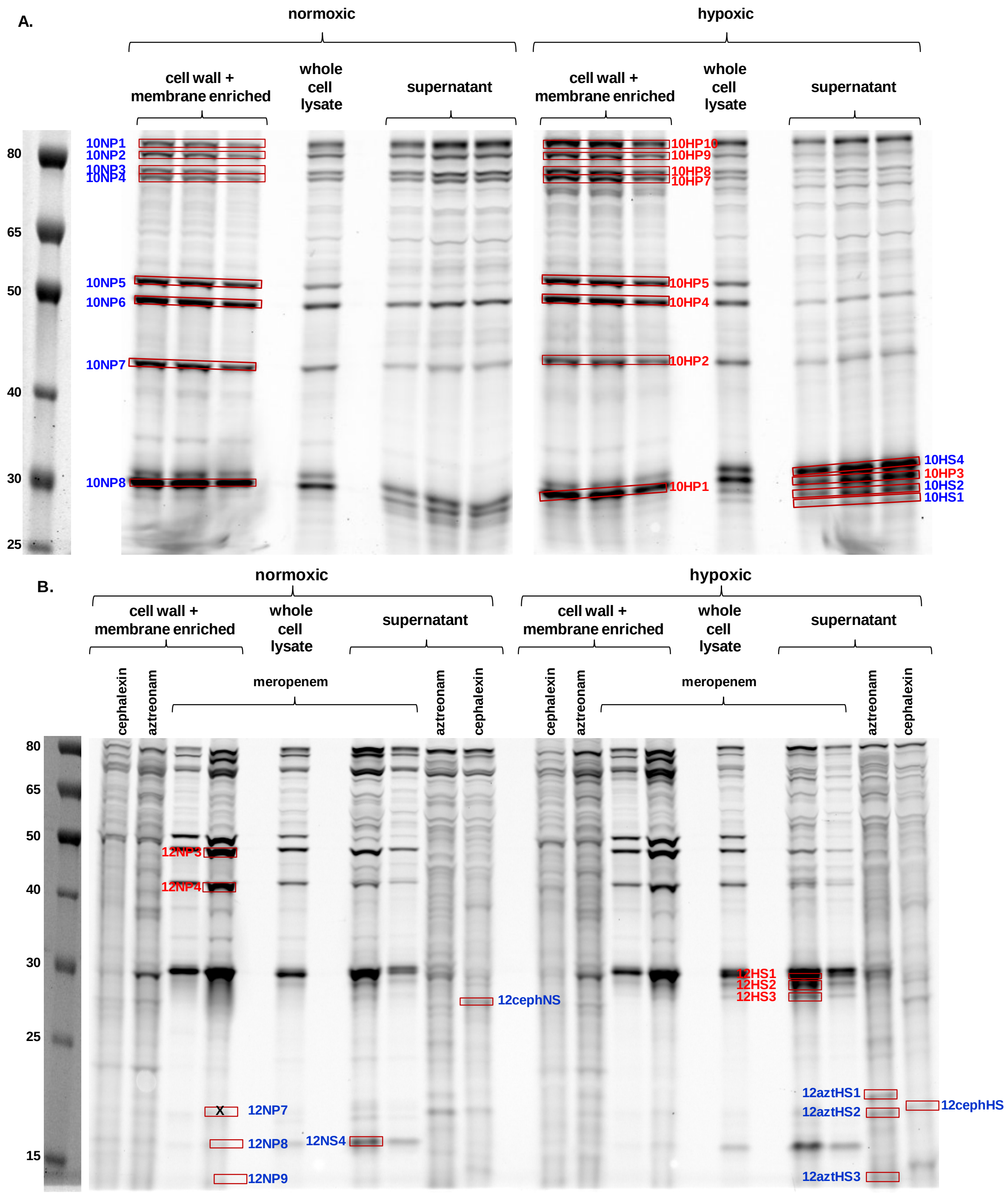


**Figure S2. Images of gels used for band excision experiments.** Bands tagged with mero-Cy5 are labeled in the format [gel %(10 or 12% Bis-tris)][growth condition (N=normoxic, H=hypoxic)][excision number]. Bands tagged with cephX-Cy5 or azt-Cy5 are labeled in the format [gel %(10 or 12% Bis-tris)][pobe (ceph=cephX-Cy5, azt=azt-Cy5)][growth condition (N=normoxic, H=hypoxic)][excision number]. Bands labeled in blue are those processed and analyzed following *Procedure I*. Bands labeled in red are those processed and analyzed following *Procedure II*. Samples from *Mtb* mc$^{2}$6020 were grown under either normoxic or hypoxic conditions and were fractionated by centrifugation into cell wall + membrane enriched and supernatant fractions[4]. **A.** All samples were labeled with 5 μM of Mero-Cy5 and separated on a 10% gel. For each sample, 25 μg of protein was resolved in two lanes of 10 μg and one of 5 μg.  All three lands were excised for identification. **B.** Samples were labeled with the indicated probe (5 μM) and separated on a 12% gel. For each sample, 20 μg of total protein was loaded per lane used for band excision.

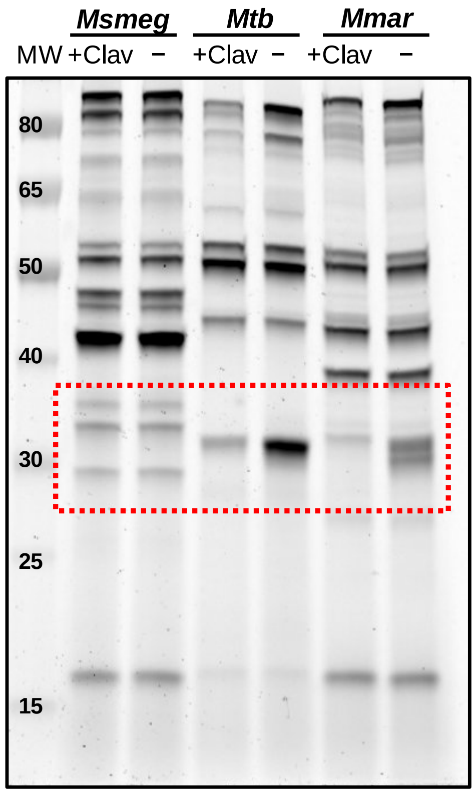


**Figure S3. Analysis of lysates from *M. smegmatis* (*Msmeg*), M. tuberculosis (*Mtb*), and M. marinum (*Mmar*).** Lysates from log-phase cultures were lysed and treated with Mero-Cy5. Lanes denoted “+Clav” were pre-treated with clavulanate (15 min) before labeling with Mero-Cy5 for 1 h. The boxed region highlights the 30 kDa region where a clavulanate-sensitive band was observed in *Mtb* and *Mmar* lysates.

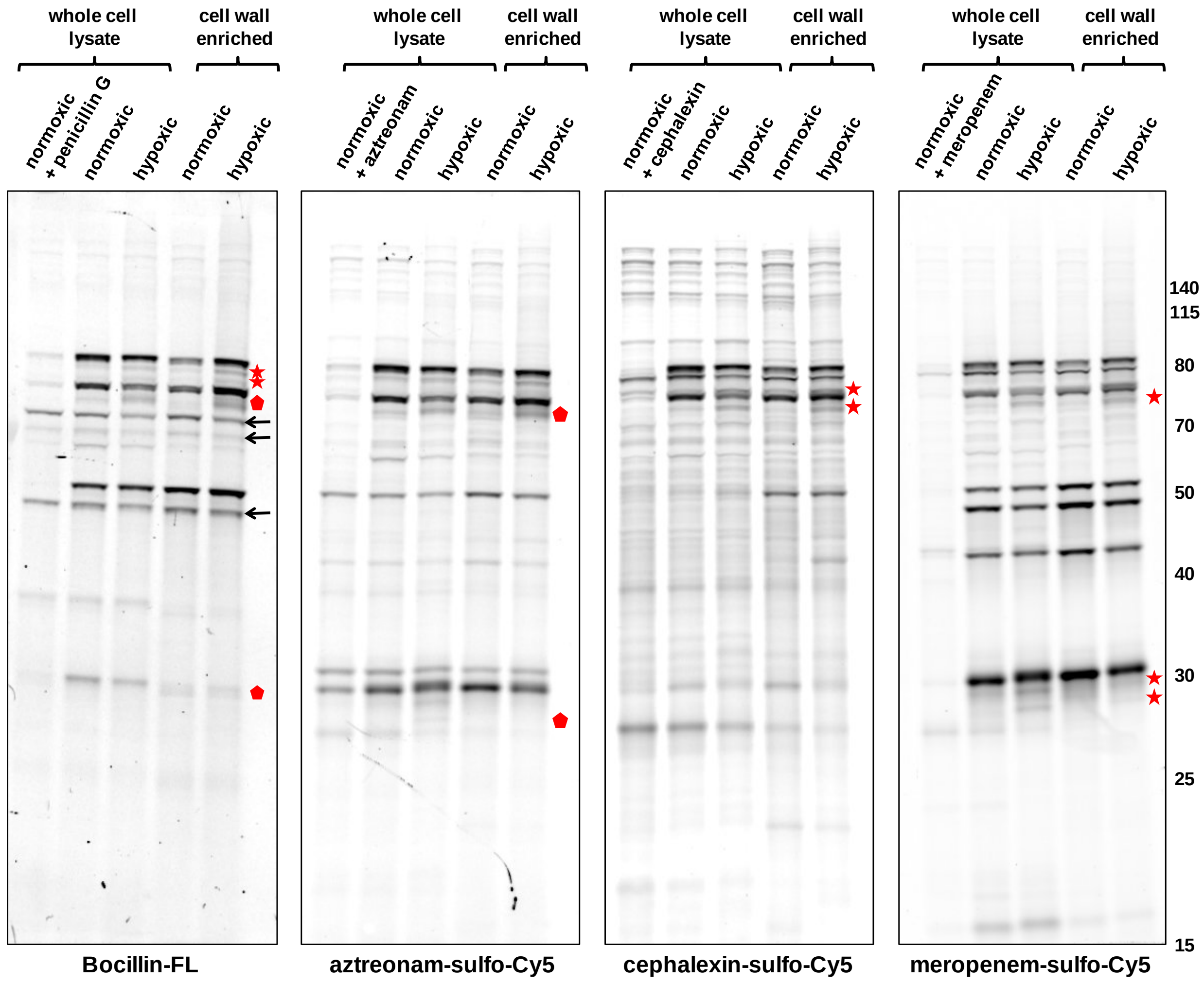


**Figure S4. β-lactam probes show differences in transpeptidase activity between actively growing and dormant *Mtb*.** Whole cell lysates of *Mtb* mc$^2$6020 grown under normoxic or hypoxic conditions were fractionated by centrifugation to enrich for cell-wall-associated material. Antibiotic pre-treatment (250 μM) occurred for 15 min at room temperature before addition of the indicated probe (**5** μM). The reactions were incubated for 80 min. Reactions were quenched by the addition of loading dye, boiled, and separated by SDS-PAGE (12% Bis-tris gel with XT-MOPS running buffer). The gel was washed and fixed overnight (20% MeOH/10% AcOH/70% $H_2O$) to remove excess probe. For Bocillin-FL the gel image was taken at $\lambda_{ex}/\lambda_{em}$ = 488/520BP40 (nm). For the other probes, Cy5 fluorescence was detected at $\lambda_{ex}/\lambda_{em}$ = 633/670BP30 (nm). ← indicates an autofluorescent band in the Bocillin-FL samples. ★ indicates a band that appears to be upregulated in dormant samples. ⬟ indicates a band that appears to be unique to dormant samples.

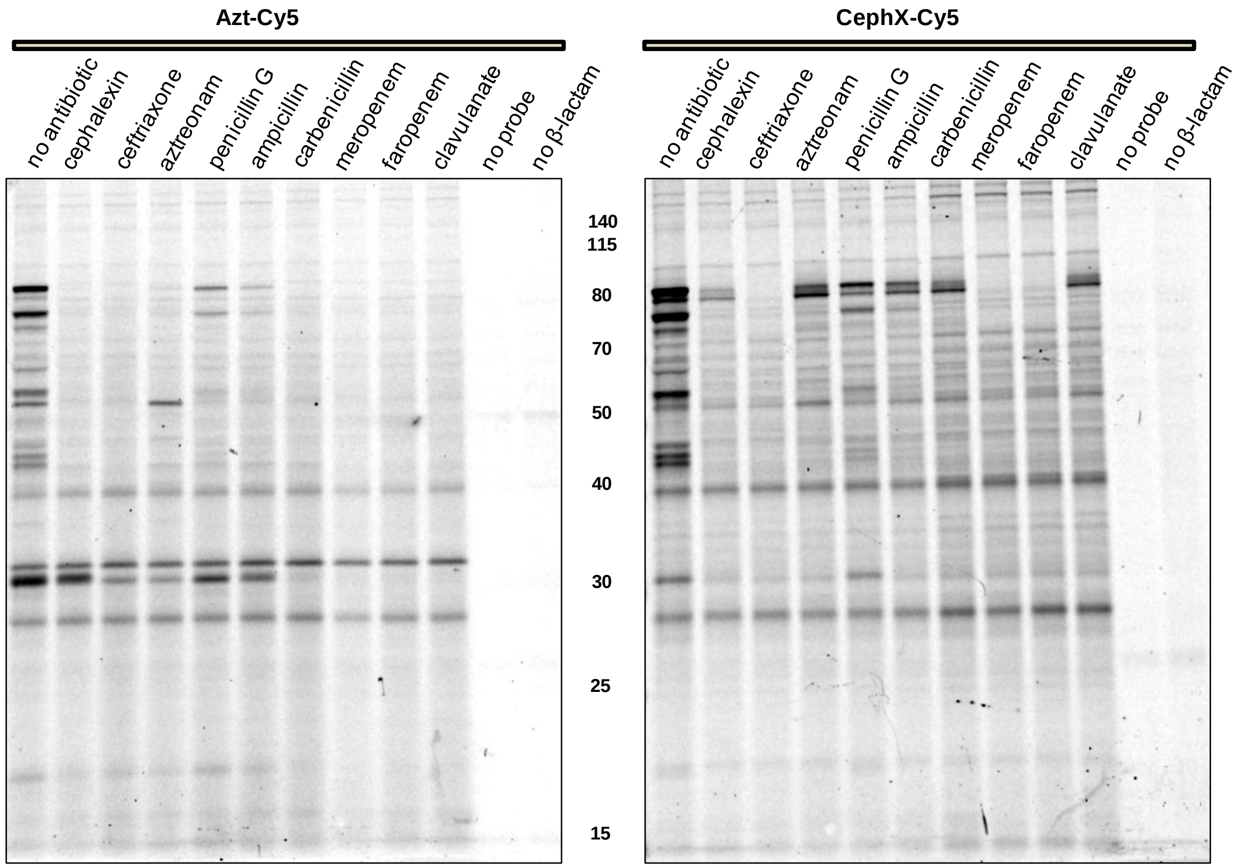


**Figure S5. Azt-Cy5 and CephX-Cy5 selectively label proteins in *Mtb* lysates.** Whole cell lysates from *Mtb* mc$^2$6020 were treated with the indicated antibiotic (500 μM) for 15 min at room temperature. The indicated probe (5 μM) was added to all samples except the "no probe" control. The reactions were incubated for 1 h. Reactions were quenched by the addition of loading dye, boiled, and separated by SDS-PAGE (12% Bis-tris gel with XT-MOPS running buffer). The gel was washed and fixed overnight (20% MeOH/10% AcOH/70% $H_2O$) to remove excess probe. Gel image was acquired at $\lambda_{ex}/\lambda_{em}$ = 633/670BP30 (nm) on a Typhoon Multimode Imager. Treatment conditions are listed above each lane. The “no β-lactam lane” was treated with a Cy5-(triazole)-butanoic acid as a control for the presence of Cy5.

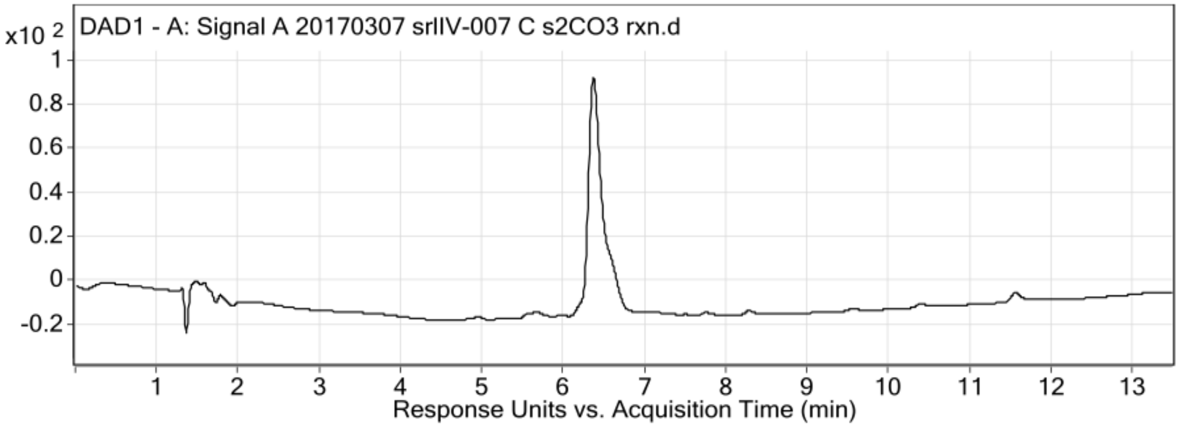


| Fragmentor Voltage | Collision Energy | Ionization Mode |
|---|---|---|
| 100 | 0 | ESI |

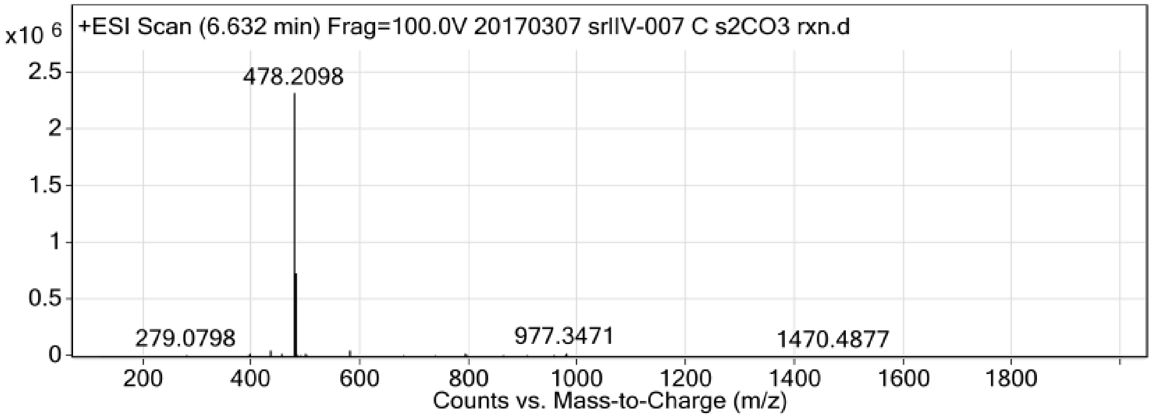


**Peak List**

| *m/z* | z | Abund |
|---|---|---|
| 396.7883 | 2 | 32040.68 |
| 434.1953 | 1 | 57403.15 |
| 478.2098 | | 2334764.49 |
| 479.1937 | 1 | 737941.13 |
| 480.1906 | 1 | 249379.38 |
| 481.1894 | 1 | 45651.85 |
| 497.1586 | 2 | 32290.44 |
| 579.3038 | 1 | 59945.37 |
| 792.5736 | 1 | 33318.93 |
| 977.3471 | 1 | 33194.94 |

**Figure S6.** LC trace and electrospray ionization results for meropenem-alkyne.

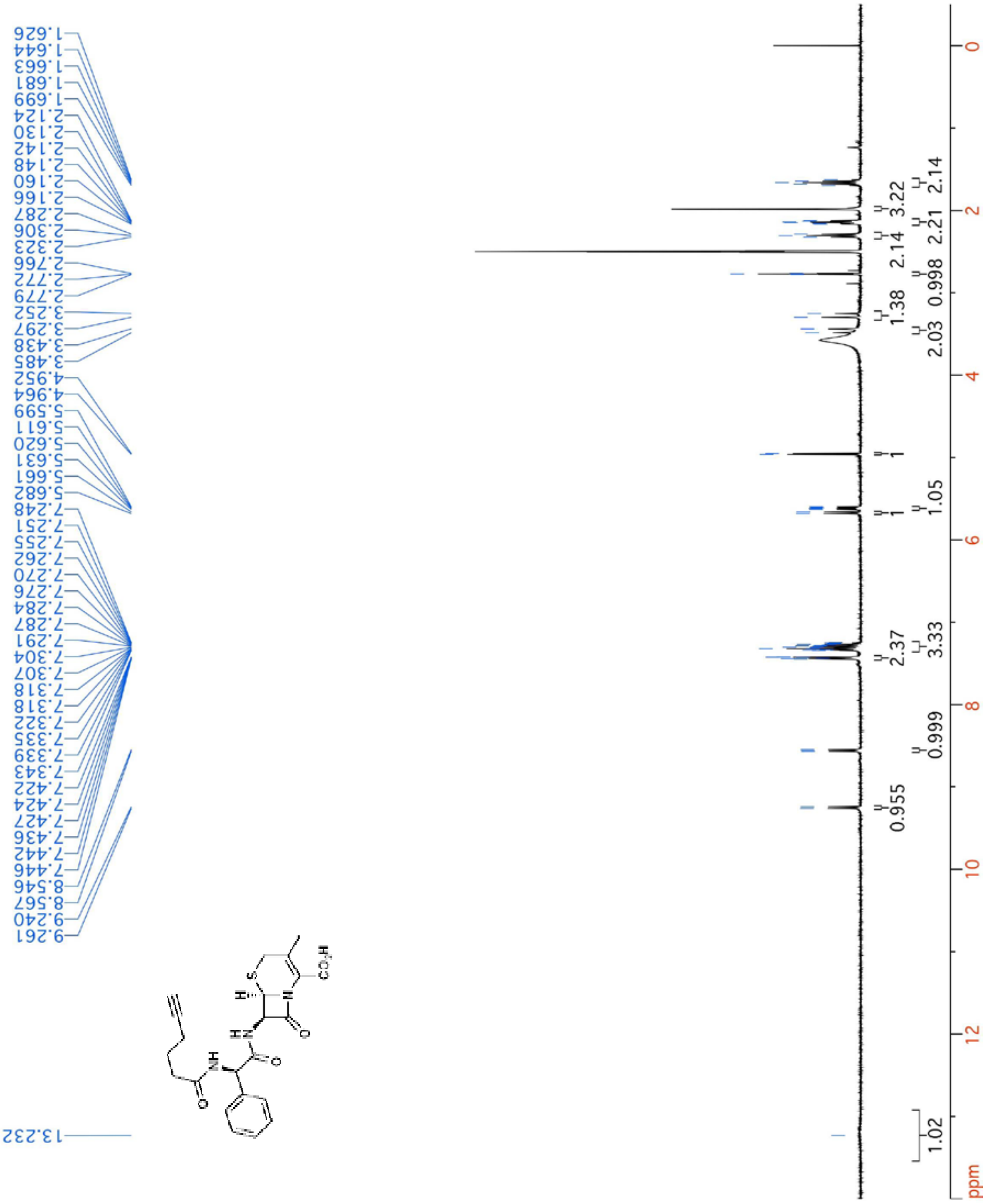


**Figure S7.** $^{1}$H-NMR of cephalexin-N-alkyne [CephX-alkyne] (400 MHz; DMSO-$d_6$).
